# Towards inferring causal gene regulatory networks from single cell expression Measurements

**DOI:** 10.1101/426981

**Authors:** Xiaojie Qiu, Arman Rahimzamani, Li Wang, Qi Mao, Timothy Durham, José L McFaline-Figueroa, Lauren Saunders, Cole Trapnell, Sreeram Kannan

## Abstract

Single-cell transcriptome sequencing now routinely samples thousands of cells, potentially providing enough data to reconstruct causal gene regulatory networks from observational data. Here, we present Scribe, a toolkit for detecting and visualizing causal regulatory interactions between genes and explore the potential for single-cell experiments to power network reconstruction. Scribe employs *Restricted Directed Information* to determine causality by estimating the strength of information transferred from a potential regulator to its downstream target. We apply Scribe and other leading approaches for causal network reconstruction to several types of single-cell measurements and show that there is a dramatic drop in performance for "pseudotime” ordered single-cell data compared to true time series data. We demonstrate that performing causal inference requires temporal coupling between measurements. We show that methods such as “RNA velocity” restore some degree of coupling through an analysis of chromaffin cell fate commitment. These analyses therefore highlight an important shortcoming in experimental and computational methods for analyzing gene regulation at single-cell resolution and point the way towards overcoming it.

## Introduction

Most biological processes, either in development or disease progression (Faith et al., 2007a; Friedman et al., 2000a; Langfelder and Horvath, 2008a; Margolin et al., 2006; Meyer et al., 2008a), are governed by complex gene regulatory networks. In the past few decades, numerous algorithms for inferring networks from observational gene expression data (Faith et al., 2007b; Friedman et al., 2000b; Langfelder and Horvath, 2008b; Margolin et al., 2006; Meyer et al., 2008b) have been developed. However, these algorithms have not been widely adopted by experimental biologists in part because they typically require an infeasible number of independent replicate observations.

A key challenge in regulatory network inference is distinguishing upstream regulatory genes from their targets directly downstream (see Methods for a formal description of the problem). Most methods that aim to do so are predicated on the notion that changes in regulators should precede changes in their targets in time (Bar-Joseph et al., 2012). Granger causality (GC) (Granger, 1969) is a statistical hypothesis test for determining whether one time series (*X*_1_) is useful in forecasting another (*X*_2_) which has been applied to infer biological networks (Zou and Feng, 2009). However, GC assumes a linear relationship between the regulator and the target, which is violated in many biological settings (Hill et al., 2016). Convergent Cross Mapping (CCM) (Sugihara et al., 2012), a more recent technique based on state-space reconstruction (Takens, 1981) can detect pairwise non-linear interactions. However, this method is limited to *deterministic* systems, and thus may be poorly suited for many cellular processes (e.g. cell differentiation), which are inherently stochastic.

Single-cell transcriptome sequencing experiments (scRNA-seq) have attracted the attention of algorithm developers working on gene regulatory network inference for two reasons. First, scRNA-seq experiments now routine produce thousands of independent measurements may open the door to sufficiently-powered inference (Liu and Trapnell, 2016). Second, algorithms that order cells along “trajectories” that describe development or disease progress offer a tremendously high “pseudotemporal” view of gene expression kinetics (Haghverdi et al., 2016; Qiu et al., 2017a; Setty et al., 2016; Trapnell et al., 2014). The recently introduced SCENIC method (Aibar et al., 2017) combines GENIE3 (Huynh-Thu et al., 2010) with regulatory binding motif enrichment to simultaneously cluster cells and infer regulatory networks. Other studies have inferred regulatory networks from scRNA-seq data using differential equations (Matsumoto et al., 2017; Ocone et al., 2015), information measures (Chan et al., 2017), bayesian network analysis (Sanchez-Castillo et al., 2017), boolean network methods (Hamey et al., 2017) or linear regression techniques (Huynh-Thu et al., 2010; Papili Gao et al., 2017; Wei et al., 2017). However, most methods don’t explicitly leverage time-series data to identify causal interactions, and more importantly, most fail to recover the correct network even in simple settings (Babtie et al., 2017; Fiers et al., 2018).

Here, we introduce Scribe, a scalable toolkit for inferring causal regulatory networks that relies on *Restricted Directed Information* (RDI) (Rahimzamani and Kannan, 2016). In contrast to GC and CCM, Scribe learns both linear and non-linear causality in deterministic and stochastic systems. It also incorporates rigorous procedures to alleviate sampling bias and builds upon novel estimators and regularization techniques to facilitate inference of large-scale causal networks. We apply Scribe to a variety of different types of real and simulated single-cell gene expression data, including single-cell RNA-seq and live imaging. In concordance with theory, we demonstrate that Scribe has superior performance compared to existing methods when the observations consist of true time-series data. However, current scRNA-seq protocols do not generate true time-series data since a cell needs to be lysed in order to sequence the transcriptome. Individual cells cannot be followed over time, breaking temporal coupling between measurements. We show a surprising result that there is a dramatic drop in performance in causal network accuracy when temporal coupling between measurements is lost. We then demonstrate that “RNA velocity”, a recently developed analytic technique for single-cell RNA-seq analysis, restores temporal coupling and improves causal regulatory network inference. Our results suggest that preserving this coupling should be a major objective of the next generation of single-cell measurement technologies.

## Results

### Scribe, a toolkit for inferring and visualizing causal regulations from single cell time-series datasets.

We aimed to develop a method for causal regulatory inference that exploits the power and resolution of single-cell RNA-seq experiments. We define *causality* as the strength of information transferred from one variable, a potential regulator, to another time-delayed response variable, a potential target, where a higher score implies stronger evidence for a causal interaction and *vice versa*. Previously, we proposed RDI as a novel information metric to accurately and efficiently quantify causality (Rahimzamani and Kannan, 2016, 2017). Built upon RDI, we developed a toolkit, Scribe, that is designed for the analysis of time-series datasets, and is especially tailored for single cell-RNA-seq **(Supplementary Figure 1, Fig 1A)**.

**Figure 1:**
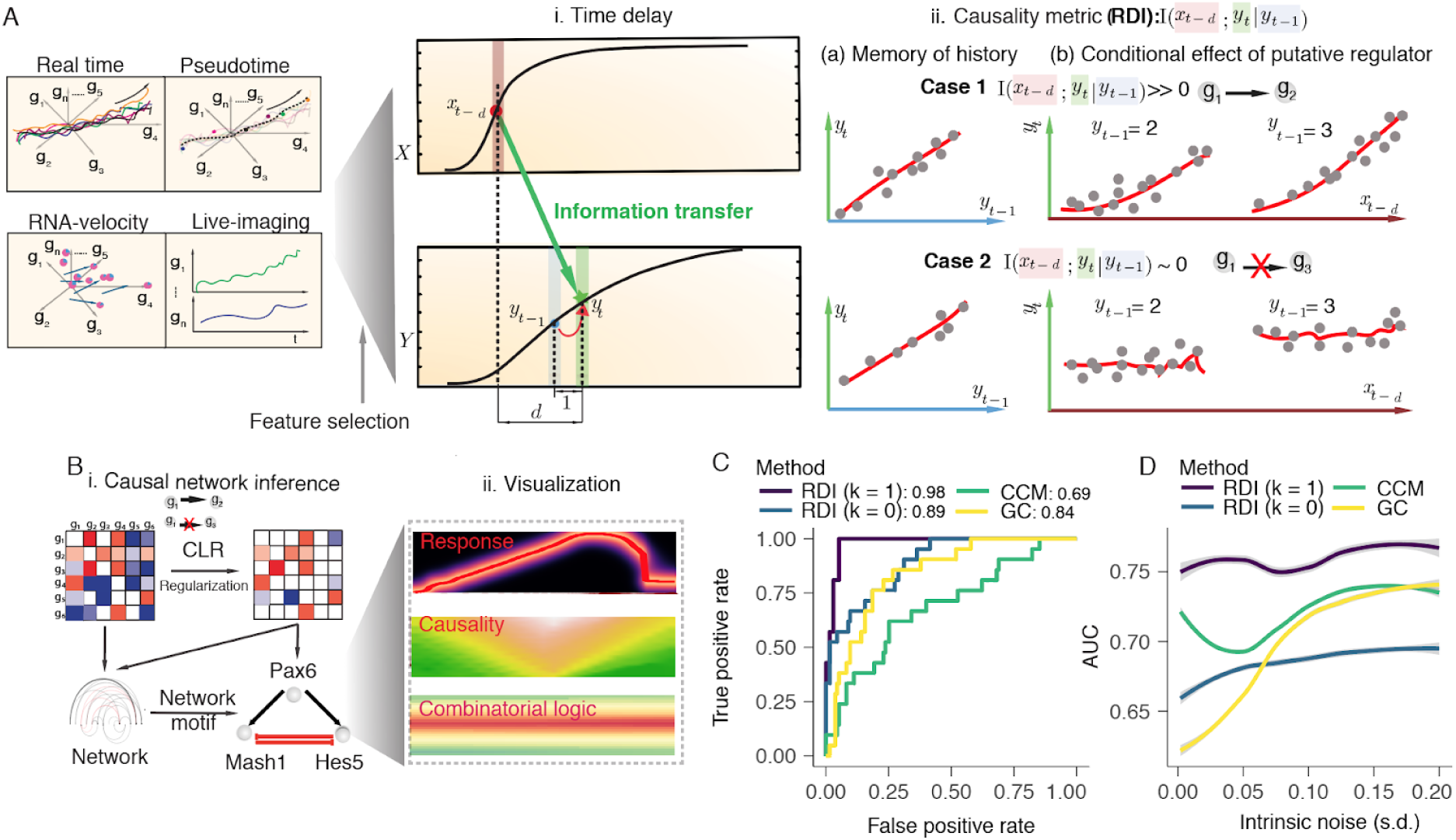
Scribe, a toolkit for inferring and visualizing causal regulations. **(A)**. Scribe detects causality from four types of single cell measurement (“pseudotime”, “live-image”, “RNA-velocity” and “real-time”) datasets with a novel information metric, restricted directed information (RDI). For a putative regulator-target pair, the current state of the target (*y_t_*) receives information from the regulator’s previous expression dynamics (*x_t-d_*) along a time-series trajectory while also having memory of its own intermediate previous state (*y_t−_*_1_). Scribe relies on RDI(Rahimzamani and Kannan, 2016) to quantify the information transferred from the potential regulator to the target under some time delay while conditioned over its past on this pseudotime-series data. A gene often has strong memory to its intermediate previous state ( *y_t−_*_1_) but RDI will only give highly positive causality score from the putative regulator to target in cases where there still is strong relationship between regulator’s history and target’s present conditioning on target’s history (Case *1 vs. Case 2*). (B) From pairwise RDI scores, Scribe assembles, refines, and prunes a directed regulatory network. Scribe also incorporates a visualization framework to visualize the response function, causal interaction as well as combinatorial regulation between gene pairs. **(C)** Receiver Operating Curves or ROC for Scribe, CCM and GC on a linear system. Standard deviation (s.d.) of independent additive noise injected to each gene at each time point and propagate through this system (intrinsic noise) is set to be 0.01. **(D)** Area Under Curve (AUC) for Scribe, CCM and GC on the non-linear neurogenesis system under different s.d. of intrinsic independent additive noise. See **Methods** on details of the simulation setup. Note that for panels **C/D**, *k* for RDI represents the number of genes to be conditioned for removing indirect causal interactions which is necessary to recover the true causal graph (Rahimzamani and Kannan 2016).

Scribe aims to be agnostic to the particular measurement technology used in an experiment, simply requiring as input time-ordered gene expression profiles for each cell as they progress along a time axis. It estimates causality scores using RDI for pairs of genes which are in turn used to build a causal regulatory network **(Fig 1A, B)**. Specifically, the RDI between two genes is formulated and quantified as the mutual information of the regulator’s past state (*x_t-d_*) and the target’s current state (*y_t_*) conditioned over the target’s history (*y_t−_*_1_) (or formally, *I*(*x_t-d_*|*y_t−_*_1_)) **(Fig 1A)**. Scribe can calculate RDI in several different ways, each of which is designed to address challenges posed by single-cell expression data. Scribe uses a novel information estimator for RDI we recently developed to account for data sparsity (Gao et al., 2017), a common feature of single-cell genomic experiments. In order to alleviate sampling biases, for example, key transitory states are underrepresented while stationary states are overrepresented, Scribe can calculate two adjusted RDI scores: termed uniformization of (conditional) mutual information (uRDI or ucRDI, respectively) to quantify the *potential causality* (*Rahimzamani and Kannan, 2017*). These scores capture how much influence a regulator can potentially exert on target *without cognizance* to the regulator’s distribution **(Supplementary Figure 1A)**.

From pairwaise RDI scores, Scribe assembles and refines a regulatory network between genes. However, many causal interactions could be indirect, and although Scribe could remove them by computing RDI between each pair of genes conditional on other potential regulators, this greatly increases the required number of samples and its running time and is typically impractical. Scribe therefore first refines the inferred network using Context Likelihood of Relatedness (CLR)(Faith et al., 2007b) and can be further sparsified with an optional method for directed graph regularization on large networks (see **Methods** for more details). Finally, Scribe offers the user a variety of ways to visualize and plot the resulting network or explore individual regulatory interactions in more detail **(Fig 1B)**.

To benchmark Scribe’s performance in inferring causal networks from time-series data, we tested it using a simplified simulation involving “real-time” measurements, in which all genes are measured in a set of individual cells that are followed overtime. We modeled the differentiation of cells in the mammalian central nervous system with a minimal regulatory network involving 12 genes through a set of linear stochastic differential equations (SDEs) based on (Qiu et al., 2012) (See Eq. 1 in **Methods)** and generated simulated measurements **(Fig 1C)**. Because real biological systems typically include both linear and nonlinear regulatory relationships, we also altered the system of differential equations that drive our hypothetical CNS differentiation system to add non-linear effects **(Fig 1D)**. We then provided these measurements as input to Scribe and compared the accuracy of the resulting network to both GC and CCM **(Methods)**. Scribe accurately recovered the causal interactions of the true network, with an Area Under Curve (AUC) score of the Receiver Operating Characteristic (ROC) curve of 0.98, while both GC and CCM inferred less accurate networks (AUC 0.84 and 0.69, respectively) **(Fig 1C, Supplementary Figure 1B)**. The performance gap between Scribe and the other methods was maintained in the presence of increasing intrinsic noise **(Fig 1D, Supplementary Figure 1B)**. In addition, we benchmark with other algorithms reported for the DREAM challenge (Hill et al., 2016), which comprises of time-series data and find Scribe performs similarly to the reported top algorithms (cRDI AUC: 0.69 while the AUC for the top three methods from DREAM challenge are 0.735, 0.6807, 0.672).

### Scribe visualizes causal regulation and combinatorial regulatory logic

Having confirmed that Scribe is able to recover correct networks, we sought ways of visualizing causality scores to aid in hypothesis generation and guide downstream validation experiments. Furthermore, in some experiments, a regulatory network may be known and the problem of interest will to detect which portions of the network are active (Krishnaswamy et al., 2014). We therefore designed several novel ways to visualize causal regulations between a putative regulatory gene *X* (or multiple regulators) and a target gene *Y* **(Fig 2, Supplementary Figure 2)**. First, Scribe plots the expected expression of the target 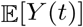 against its immediate past *Y*(*t-1*) and the expression of the regulator in the recent past *X*(*t-d*) **(Fig 2B)**. For example, in our simulated network, *Mash1* represses *Hes5:*

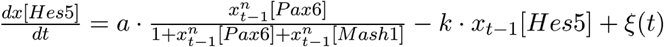

**Figure 2:**
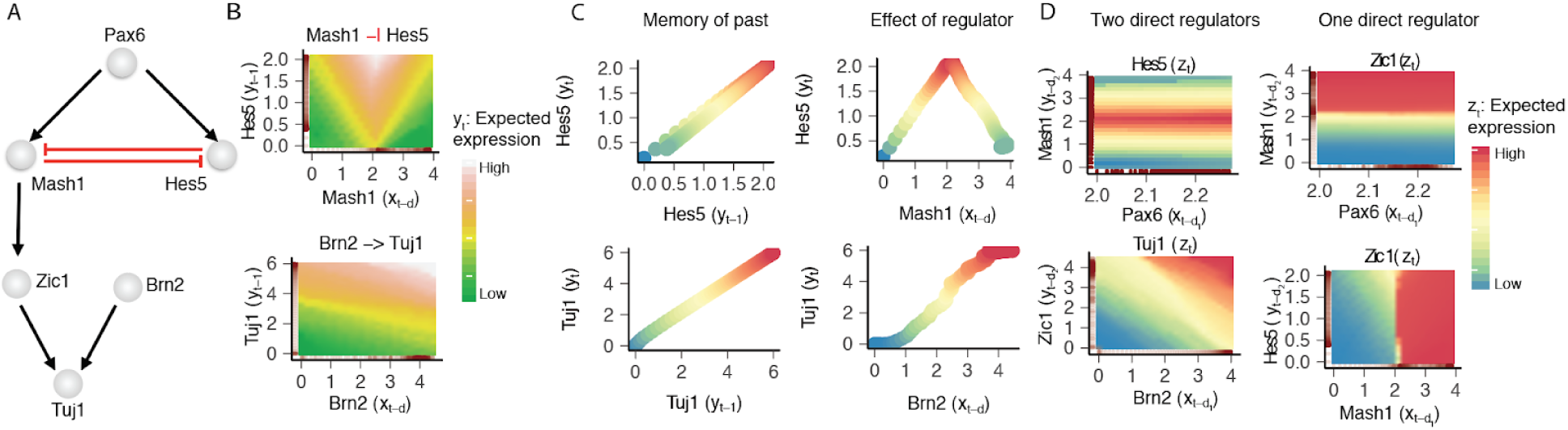
Visualize gene regulations with Scribe. **(A)** The subnetwork of regulatory interactions visualized by Scribe in panels **B-D**. See **Supplementary Figure 2A** for the full network. **(B)** A *causality* visualization reveals the information transfer from one gene to another. The horizontal axis corresponds to the regulator’s previous expression with a time lag *d* while the vertical axis corresponds to the target gene’s very recent past expression. The heatmap color scale corresponds to the expectation of the target gene’s current expression given the target’s its very recent past and the putative regulator expression with a time lag *d*, or 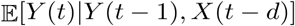 (**C**) Scatterplots describing relationships of gene regulatory interactions with time delay. **(D)** A *combinatorial regulation* visualization reveals the combinatorial gene regulation from two regulators to a target gene. X-axis corresponds to the regulator’s previous expression with a time lag *d*_1_ while y-axis corresponds to another regulator’s gene expression with another time lag *d*_2_. The heatmap corresponds to the expected value of the target gene’s current expression given both of the regulators’ expressions with the time lags (*d*_1_ or *d*_2_) or 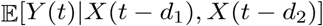 For this simulation, all the time delays *d, d*_1_ or *d*_2_ are set to be 1.

Scribe shows the relationship between these two genes to follow a pattern reminiscent of threshold inhibition **(Fig 2B,C)**.

In our hypothetical system, *Tuj1* is regulated by *Brn2*, *Zic1*, and *Myt1l* as described by the following ordinary differential equation:

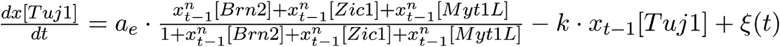

Scribe’s visualizations revealed the sigmoid activation relationship between *Brn2* and *Tuj1* **(Fig 2B, C)**. Scribe also includes a second type of visualization, showing the time-delayed response function of a target to a regulator **(Supplementary Figure 2B)**.

Most genes are regulated by more than one upstream factor and many regulatory relationships are indirect, so we developed a third and novel visualization method that helps distinguish direct and indirect regulations and dissect combinatorial regulatory logic between sets of genes. Scribe renders the expected expression of a target conditional on the space of possible expression values of two upstream regulators at some point in the recent past **(Fig 2D)**. For example, Scribe shows that *Tuj1* varies over a simple linear gradient of the cumulative levels of *Brn2* and *Zic1*, consistent with their direct, additive role on *Tuj1* levels in equation 1 above. In contrast, *Zic1* depends directly on *Mash1* and *indirectly* on *Hes5*. Scribe’s visualization of *Zic1* expression conditional on past values *of Mash1* and *Hes5* reveals very modest dependence on *Hes5*, but strong dependence on *Mash1* expression. These examples show that in principle, Scribe’s visualizations can help reveal combinatorial regulation logic and distinguish direct, causal regulatory interactions from indirect ones.

We next used these three visualizations to systematically examine all pairs of interactions between genes in the hypothetical network, which revealed them to be consistent with our prior characterization **(Supplementary Figure 2;** c.f. **Fig SI5A from** (Qiu et al., 2012)). We also systematically tested various regulatory network “motifs” of genes to confirm that these visualizations correctly reveal the causal regulatory relations between them **(Figure 3)**. Together, these analyses demonstrate that Scribe not only can recover causal regulatory interactions between genes, but also provides the user with tools to explore gene interactions in great detail, facilitating the design of follow up experiments.

**Figure 3:**
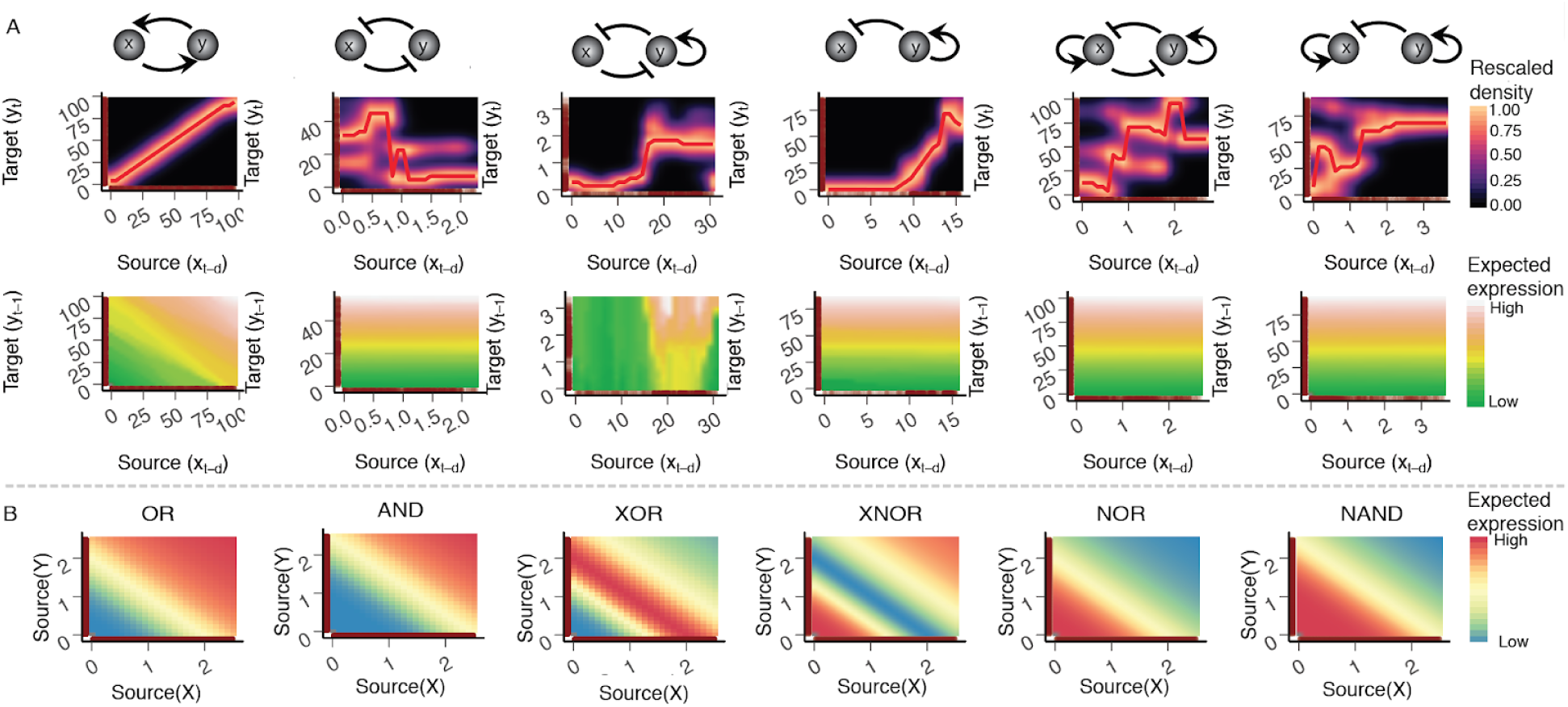
Visualizing pairwise interactions from robust two-gene network motifs and combinatorial regulations from common two-input logic gates with Scribe. **(A)** Visualizing response and causality for two-gene network motifs. Top: robust network motifs from (Ma et al., 2009); middle: corresponding response visualization plots, similar to the DREVI plot reported in (Krishnaswamy et al., 2014) with the difference that we explicitly consider the time-delayed response from the target to the regulator (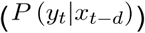) (see **methods** for more details); bottom: corresponding causality visualization plots. The first node in the motif plot corresponds to the source (x-axis) while the second the target (y-axis). **(B)** Visualizing combinatorial regulation for six two-input logic gates (OR, AND, XOR, XNOR, NOR and NAND).

### Causal network inference of *C. elegans’* early embryogenesis with “live-imaging”

In order to assess the performance of Scribe in practice, we examined *Caenorhabditis elegans’* early embryogenesis, where live-imaging has been used to measure nearly half of all transcription factors’ protein expression dynamics in every single cell in an embryo (Murray et al., 2012). This dataset consists of 265 time series that each track the expression dynamics of a transcription factor using fluorescent reporter constructs. Measurements were collected at one minute intervals in every cell of the developing embryo for the first ~350 minutes of embryogenesis **(Fig 4A)**.

**Fig 4:**
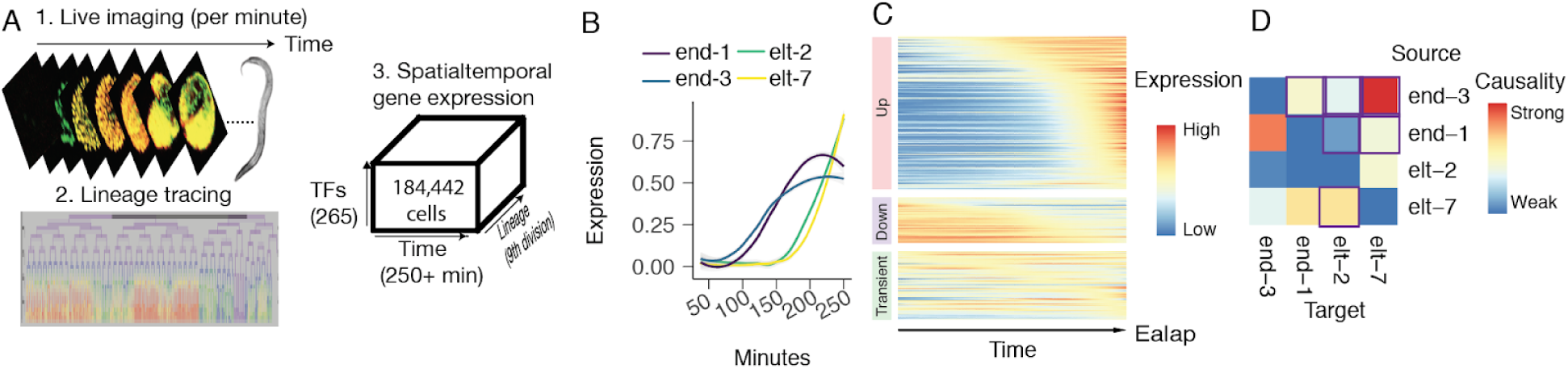
Living imaging dataset of *C. elegans’* early embryogenesis captures transcription expression dynamics hierarchy. **(A)** Scheme used by Murray *et* a/for measuring transcription factors protein expression dynamics in real-time for every cell during early *C. elegans* embryogenesis. Protein-RFP fusions reporters are used to measure transcription factor protein expression levels with 3D live imaging every minute in each cell while a ubiquitous histone-GFP marker is used to trace the *C. elegans* cell lineage. Reporter fluorescence data was then mapped onto the invariant cell lineage. Combining expression measures in each corresponding cell from each embryo yields a tensor with dimensions of 265 genes X 550 time points X 1365 (in total more than 180,000 data points, after removing those with invalid measurements). **(B)** Single cell lineage-resolved fluorescence data captures temporal dynamics of *E* lineage master regulators during C. *elegans* embryogenesis. The expression for each gene is scaled to be between 0 and 1 and then smoothed using LOESS regression. (**C**) Expression dynamics for 265 report TFs along the lineage leading to the *Ealap* cell. The entire developmental lineage from the first *E* cell all the way to the *Ealap* cell in each embryo for each TF reporter is used to make the heatmap. The raw fluorescence intensity is scaled to be between 0 and 1 and then smoothed using LOESS regression. The order of genes in each row is calculated as previously described (Pliner et al., 2017). **(D)** Scribe reconstructs the causal regulatory network for the four master regulators (*end-1/3, elt-2/7*).

We tested whether Scribe was able to learn validated genetic interactions that govern worm development. For example, in the intestinal cell lineage *Ealap* the transcription factors end-1 and end-3 were upregulated prior to their targets elt-2 and elt-7 **(Fig 4B)**, and well before most other upregulated factors in this lineage **(Fig 4C)**. We then ran Scribe on these four genes to determine whether it could correctly infer the causal regulatory interactions between them. Although Scribe captured some known causal interactions among the core transcription factors that specify this lineage (Owraghi et al., 2010), it also reported both false positive and false negative interactions. For example, Scribe reports that end-1 strongly regulates end-3 **(Fig 4D)**. Although accurate network inference surely depends on many factors, we hypothesized that the false positives and false negatives in the live imaging analysis were mainly due to the inability of the system to report measurements for more than one gene in the same individual cells. Because each reporter strain tracks a different gene (in a different animal), fluctuations in expression levels of a regulator are not “coupled” to corresponding fluctuations in its targets.

### Accurate causal network inference requires temporally coupled expression data

Next, we explored Scribe’s ability to recover causal interactions using single-cell RNA-seq which in contrast to live-imaging measures many genes in each cell. We first collected publically available datasets from several biological systems including developing airway epithelium (Treutlein et al., 2014), dendritic cell response to antigen stimulation (Shalek et al., 2014), and myelopoiesis (Olsson et al., 2016). We then pseudo-temporally ordered these cells as previously described using Monocle 2 (Qiu et al., 2017a). Next, we ran Scribe on these pseudotime series **(Fig 5, Supplementary Figures 3, 4, 5)** and examined the regulatory interactions reported for known transcriptional regulators of these systems. For each gene, we summed the causal interaction scores to all other genes, deriving a measure of its aggregate influence on the system. Reassuringly, these aggregate causality scores were significantly higher for known transcriptional regulators than for genes believed to be targets by the authors of the original studies **(Fig 5)**. Moreover, Scribe identified several regulatory interactions, such as *Gata1-Gfi1-Klf4*, which are known to play an important role in myeolopoeisis (Laslo et al., 2006; Stopka et al., 2005; Tamura et al., 2015) **Supplementary Figure 5I)**. In recovering known regulatory interactions in each system, Scribe marginally outperformed GC and CCM but all three methods generally performed poorly, with no method reaching an AUC of greater than 0.7 **(Supplementary Figure 5)**.

**Figure 5:**
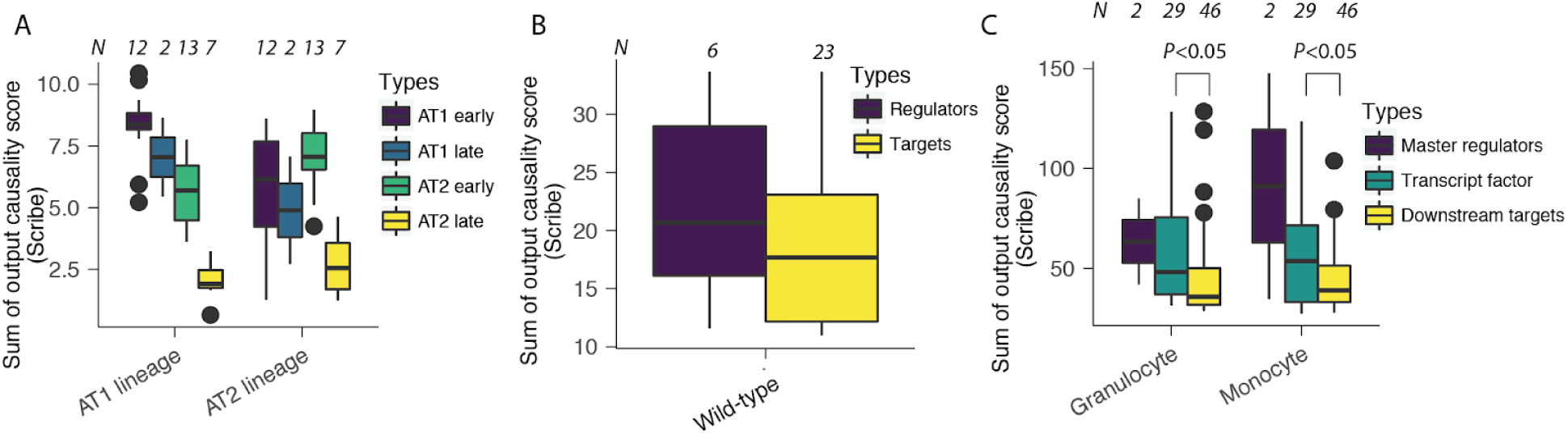
Scribe correctly reveals the ordering of the sum of outgoing RDI for a variety of single-cell RNA-seq datasets. In order to demonstrate Scribe’s power in detecting the direction of causal regulation, we test the hypothesis that the sum of outgoing edges’ causality score should be higher in groups of potential regulators than in their targets on three different datasets: lung (Treutlein et al., 2014), LPS (Shalek et al., 2014) and myelopoeisis (Olsson et al., 2016). **(A)** Total outgoing causality scores of the putative regulators is higher compared to that of the target genes across AT 1 or AT2 branch. **(B)** Same as in panel A but for the LPS data (only wild-type cells are chosen from this dataset to avoid testing on disrupted LPS response network in the knockout cells). (**C**) The master regulators have the highest total outgoing causality scores compared to the putative direct targets (transcription factors) and then the putative secondary targets (downstream targets). To obtain total outgoing causal scores, causal scores between all gene pairs are calculated with RDI and then processed by the CLR algorithm, followed by summing up all outgoing edges’ scores for each gene. Integers (N) above each boxplot corresponds to the number of genes used for creating the plot. An unpaired two-sample t-test is used to test each pair of hierarchical groups of genes. Only pairs of genes detected as significantly different (p < 0.05), which also happen to include larger number of genes, are labelled.

We hypothesized that as with live imaging datasets, lack of coupling between the expression measurements in pseudo-temporally ordered single-cell RNA-seq data leads to poor accuracy during regulatory network inference. Although single-cell RNA-seq measures many genes, each cell is sampled (destructively) only once. In contrast to true time series in which an individual cell is followed and measured longitudinally, in pseudo-temporal datasets, each expression measurement comes from a different cell. Therefore, the gene expression fluctuations of a regulator do not propagate to its target at some later point in pseudotime.

To test whether causal network inference requires temporal coupling between genes across measurements, we ran Scribe on simulated data collected using four strategies for obtaining longitudinal measurements from individual cells. First, we consider “real-time”, an ideal theoretical technology in which all genes are tracked in each individual cell as that cell differentiates. Current single-cell techniques are limited either to the measurement of just a handful of genes over time (e.g. live imaging of fluorescent reporters), or require destroying the cell to collect data (e.g. sc-RNA-seq). We therefore consider a second setting “live-imaging”, in which each cell is tracked over time but only one gene is measured. Third, we examine pseudotime, where all genes are measured only once in distinct cells that have been sampled from a population undergoing differentiation. Finally, we tested Scribe on RNA velocity data, which consists of a snapshot measurement of each cell’s current transcriptome along with a prediction of that same cell’s expression levels at a short time in the future **(Fig 6A)**.

**Figure 6.**
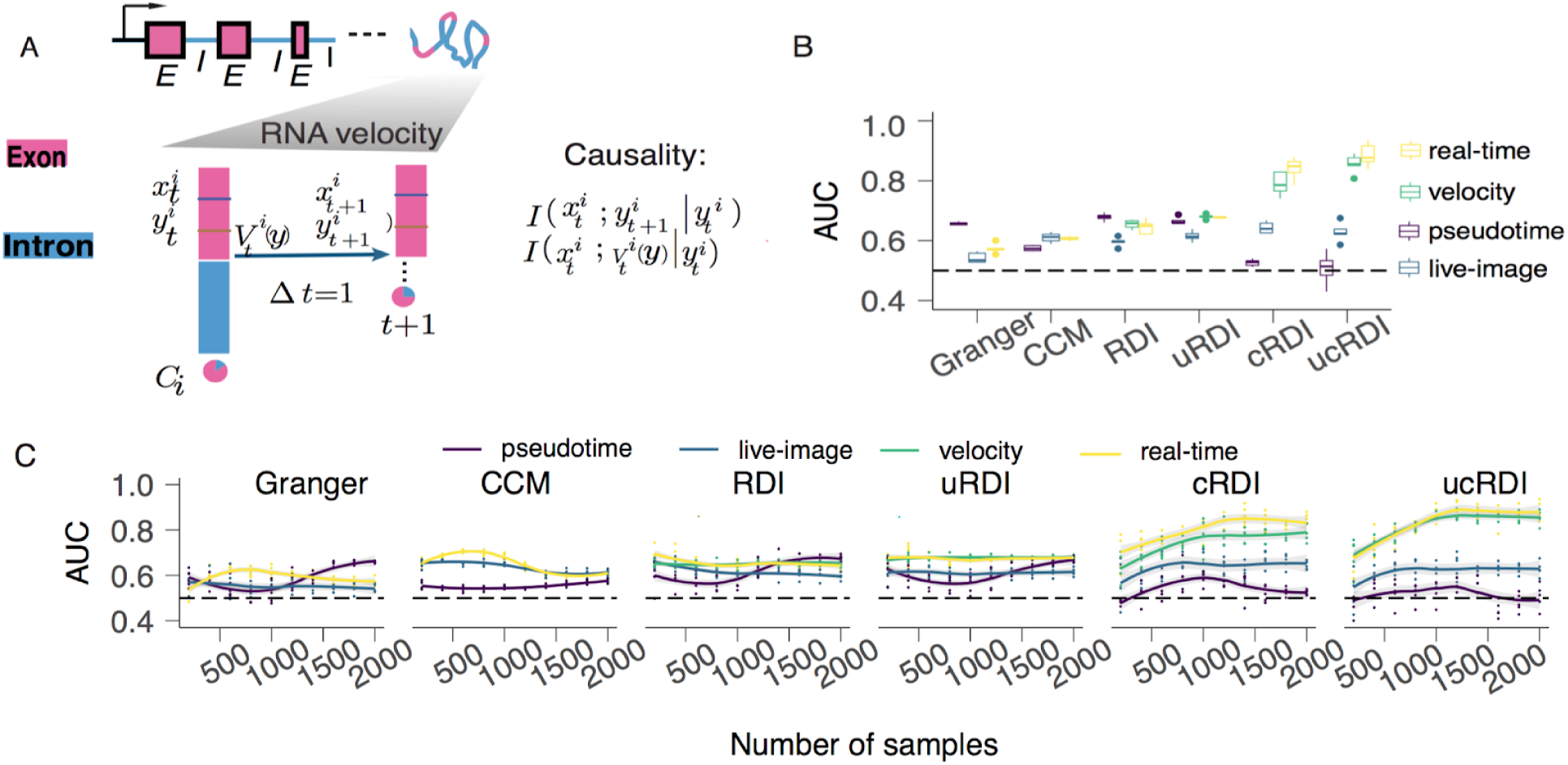
Temporally coupled gene expression measurements are necessary for inferring causal regulatory interactions. Four possible types of single-cell time-series datasets, pseudotime (no dynamics coupling across genes and time-points), live-imaging (dynamics coupled for each gene across time-points but not across genes), “RNA-velocity” (dynamics coupled for all gene between two time points measured in each cell) and real-time (dynamics coupled for all gene across all time-points in each cell), are simulated (see **Methods** for details). **(A)** Incorporating RNA velocity analysis into Scribe for causal inference. A gene with multiple exons (pink box, ***E***) and introns (blue line, ***I***) is transcribed into immature RNA and then spliced into mature RNA, both of which can be quantified by scRNA-seq. See **Methods** section *Benchmarking Scribe with alternative algorithms on inferring causal regulatory network*. (B) Dynamics coupling in real time and “RNA velocity” datasets enables Scribe to correctly infer causal regulatory network. Scribe (including four variants, RDI (*k* = 0), uRDI ( *k =* 0), RDI (*k* = 1) and uRDI (*k* = 1)) and two alternative causal inference methods, Granger and CCM are used to reconstruct network based on 2,000 data points from each of the four different single-cell time-series datasets. The results from each method are then compared with the known network architecture to obtain the AUC score. Five replicates are performed for each methods. **(C)** Dependence of causal inference on the number of samples. The same analysis as in **A** is performed but with downsampled datasets on a sequence from 200 to 2000, incremented by 200, data points (see **Methods** for more details). RDI: restricted directed information. uRDI: uniform RDI which replaces the biased data distribution with a uniform distribution to remove the dependence of the sample distribution.

Using pseudo-temporal measurements, Granger causality, convergent cross mapping, and Scribe all performed very poorly in recovering direct, causal interactions between genes in the hypothetical network **(Fig 6B, Supplementary Figure 6A)**. The inability of these methods to recover regulatory interactions is unlikely to be due to undersampling of the system, as performance was insensitive to varying the number of cells captured in the simulated datasets **(Fig 6C, Supplementary Figure 6B)**. Performance of the three methods was only modestly better when using data captured by “live imaging”, in which one gene is followed in each cell over time, but no two genes are ever measured in the same individual cells.

We next evaluated two alternative modes of measuring gene expression dynamics in single cells in which fluctuations are coupled. Using conditional Restricted Directed Information, Scribe produced highly accurate reconstructions from “real time” measurements of gene expression (AUC: 0.859 ± 0.0283), in which every gene is measured repeatedly in a set of cells as they differentiate. This demonstrates that when measurements are fully coupled across time, and fluctuations in a regulator can propagate to its targets, restricted directed information correctly reveals causal regulatory interactions. RNA-velocity offers the ability to perform causal inference via RDI based on two data points from the *same* cell **(Fig 6A)**. Encouragingly, Scribe also recovered accurate networks (AUC: 0.837 ± 0.0189) with “RNA velocity” measurements. Although RNA velocity does not repeatedly measure cells, it provides a “prediction” of the future expression levels of each gene based on comparing mature to immature transcript levels, in effect introducing a form of temporal coupling to the data. These simulations show that methods for regulatory inference based on information transfer fail using data from measurement modalities in which fluctuation of a regulator’s expression across cells is “uncoupled” from fluctuations in it’s targets.

### Causal network inference with “RNA-velocity” reveals regulatory interactions that drive chromaffin cell differentiation

We next sought to test whether Scribe could recover causal network interactions using real RNA velocity measurements. Recently, La Manno and colleagues applied RNA-velocity to study the chromaffin cells differentiation as well as their associated cell cycle dynamics (La Manno et al., 2018). We used this chromaffin dataset as a proof-of-principle for incorporating “RNA velocity” into Scribe. We first reconstructed a developmental trajectory from mature mRNA expression levels from each cell in this dataset and then applied BEAM (Qiu et al., 2017b) to identify genes that significantly bifurcate between Schwann and chromaffin cell branches **(Fig 4B)**. Reassuringly, these genes were enriched in processes related to neuron differentiation along the path from SCPs (Schwann Cell Progenitors) to mature chromaffin cells **(Supplementary Figure 7A)**.

We then applied Scribe to the RNA velocity measurements from the 3,665 significantly branch-dependent genes (qval < 0.01, Benjamini-Hochberg correction) **(Figure 4C, Supplementary Figure 7)**. We first built a network between significant branching transcription factors (TFs) as well as from TFs to the significant targets in chromaffin lineage and found that only 0.75% of TFs interact with each other while 8.40% TFs regulate potential targets (causality score > 0.05) **(Supplementary Figure 7D)**. We then inferred a core network between fourteen TFs believed to drive chromaffin cell differentiation (Furlan et al., 2017). Within this core network, Scribe identified two feed-forward loop (FFL) motifs (Alon, 2007); *Eya1-Phox2a-Erbb3* and *Gata3-Phox2a-Notch1* **(Fig 7C-E)**. The STRING database of genetic and molecular interactions (Szklarczyk et al., 2017) provided additional support for these regulatory motifs **(Supplementary Figure 7D)**. These network motifs were not found by Scribe when run on pseudo-temporally ordered, mature mRNA measurements alone (data not shown).

**Fig 7:**
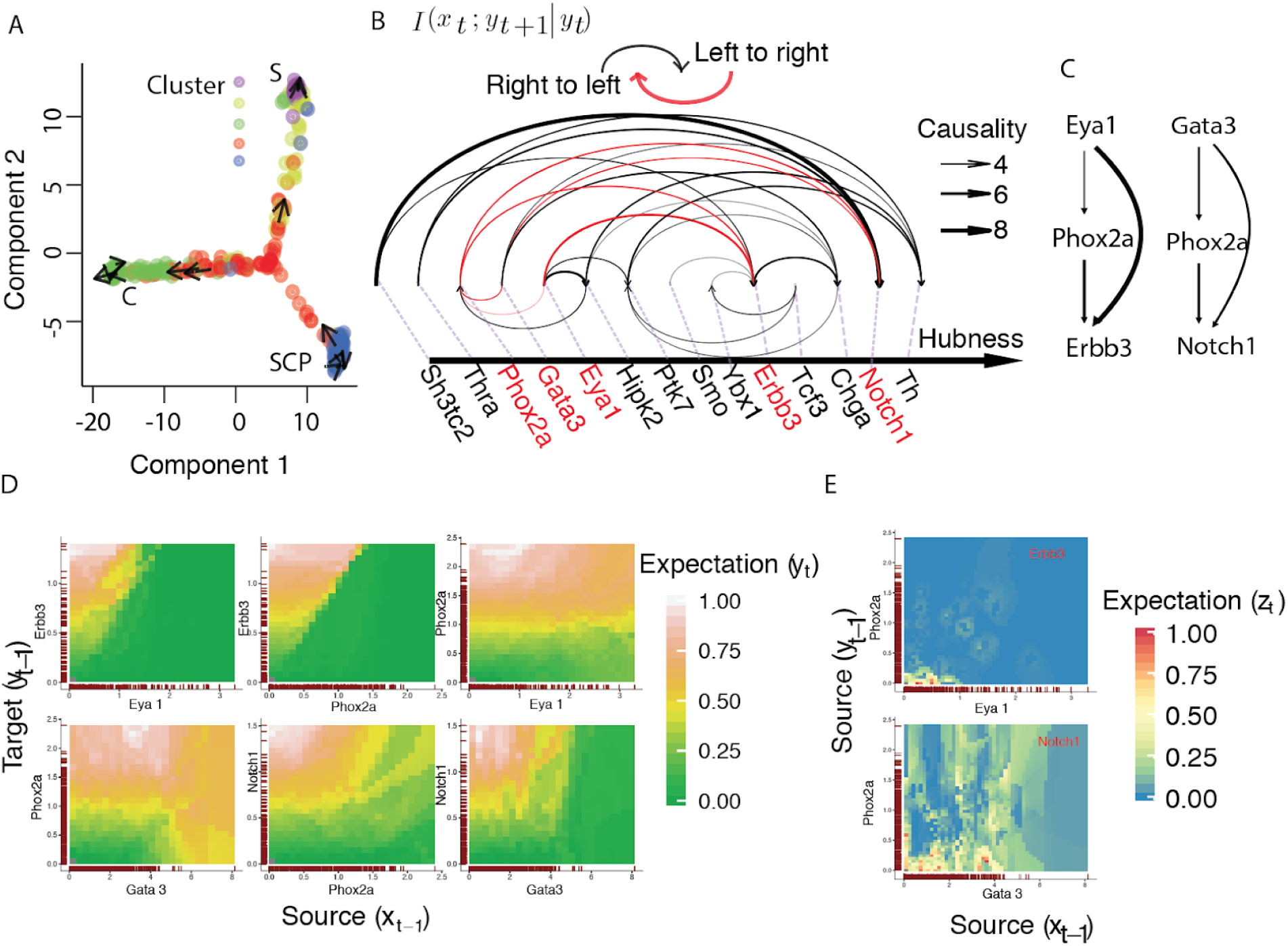
Causal inference in Scribe with RNA-velocity. **(A)** RNA-velocity vector projected onto the first two latent dimensions. A small subset of arrows are used to visualize the velocity field of the cells. **S:** Sympathoblasts; **C:** Chromaffin. **SCP:** Schwann Cell Progenitor. The color of each cell corresponds to the cluster id from **Fig 5B** of ref. (Furlan et al., **2017)**. Only the exon values from RNA-velocity framework are used to reconstruct the developmental trajectory. All **384** cells used in (La Manno et al., **2018)** are used (same as below). **(B)** A core causal network for chromaffin cell commitment inferred based on RNA-velocity. Gene set is collected from ref. (Furlan et al., **2017)**. CLR (context likelihood of relatedness) is used to remove spurious causal edges (quantified with *i*{*xl\vi+i\vl*)) in the network (see **methods)**. Network is layouted as an arc-plot where all the genes are ordered on a line, sorted by the hub centrality score (see **Methods)** decreasing from left to right. The edges above (below) the line indicate the interactions from the left (right) genes to the right (left). The width of the edge corresponds to the normalized causality score returned after applying CLR on RDI values. Genes are labeled below the horizontal line. **(C)** Two potential coherent FFL (feed-forward loop) motifs of chromaffin differentiation are discovered from the core network. Edge width corresponds to causal regulation strength. **(D)** Visualization of the six causal regulations pairs in the feedforward loops of *Eya1-Phox2a-Erbb3* and *Gata3-Phox2a-Notch1*. Current mature mRNA and predicted future mRNA estimated from “RNA velocity” analysis framework are used as input and smoothed using as a local average. Expected values are then rasterized to plot as a two-dimensional heatmap (See methods for details). (**E**) Visualizing combinatorial regulation logic for the two feedforward loops in Panel **C** with Scribe. For both Panels ***D*** and **E**, a grid with 625 cells (25 on each dimension) is used. Similarly, expectation values are scaled by the maximum to obtain a range from 0 to 1.

## Discussion

Causal gene regulatory network inference requires a large volume of data and is vastly easier in the context of time series experiments. Single-cell RNA-seq experiments can produce thousands of observations which when pseudo-temporally ordered reveal gene expression dynamics at extraordinarily high resolution. The technology has understandably sparked renewed interest in network inference algorithm development. Recently, methods to leverage single-cell data for network inference based on mutual information and pseudotime ordering have been reported (Chan et al., 2017; Hamey et al., 2017; Huynh-Thu et al., 2010; Matsumoto et al., 2017; Ocone et al., 2015; Papili Gao et al., 2017; Sanchez-Castillo et al., 2017; Wei et al., 2017); however, most of those methods only report statistical dependence (Chan et al., 2017; Huynh-Thu et al., 2010; Papili Gao et al., 2017; Sanchez-Castillo et al., 2017), and as shown by Matsumoto *et al*, many return networks only slightly more accurate than random guessing (Matsumoto et al., 2017). Despite extensive research into gene regulatory network inference over the past several decades, the fundamental source of poor performance by these methods on single-cell data remains uncertain. One possibility is that, even with the tremendous gains in throughput acheived by developers of single-cell RNA-seq technology over the past decade (Svensson and Vento-Tormo, 2017), these methods still haven’t been provided with enough data to accurately reconstruct networks. Alternatively, the basic approach of inferring genetic interactions based on statistical interactions between their measured expression levels may be fundamentally limited.

We developed Scribe, which uses recently reported advances in information theory to infer complex *casual* regulatory interactions between genes. Scribe employs Restricted Directed Information (RDI), overcoming limitations inherent to Granger Causality (GC) and Convergent Cross Mapping (CCM). Scribe also provides several ways to visualize causal information transfer, helping users distinguish between direct and indirect interactions and unravel combinatorial regulatory logic.

Although Scribe correctly infers causal regulatory interactions in simulated measurements that track all genes in an individual cell over time, it performs poorly on live imaging or pseudo-temporally ordered single-cell datasets. We demonstrate that poor performance is due to the loss of temporal coupling between measurements of genes that interact, in which fluctuations in levels of a regulator propagate to measurements of its targets. This may explain poor performance by a broad class of information theoretic or statistical approaches for inferring regulatory networks from single-cell RNA-seq data. If so, then simply improving the throughput of single-cell RNA-seq protocols will not be sufficient to power inference methods.

Improvements to single-cell expression assays that produce measurements for multiple genes that are coupled across time may enabled accurate regulatory network inference using Scribe or similar approaches. Although methods for nondestructively tracking expression levels of many genes in single cells over time have not been described, several assays have been reported that provide snapshot estimates of both steady state mRNA levels along with their rates of synthesis. These assays report measurements of the current and future transcriptome of individual cells, essentially providing temporal coupling over a short time horizon. For example, SLAM-seq (Herzog et al., 2017; Muhar et al., 2018) orTUC-seq (Riml et al., 2017) assay mature RNA levels and estimate the rate of their synthesis via nucleotide labeling or conversion based approaches. Sequential, multiplex RNA FISH or “Seq-FISH” (Shah et al., 2018), probes both exons and introns of RNAs can also provide similar measurements. RNA velocity, which analyzes single-cell RNA-seq reads falling within introns, estimates both mature mRNA levels and their immature intermediates to predict the transcriptome a short time in the future, also generates coupled measurements. Accordingly, using RNA velocity measurements greatly improves Scribe’s accuracy compared running it on pseudo-temporal single-cell RNA-seq measurements.

Gene regulatory network inference from observational measurements of gene expression is widely regarded as amongst the most difficult problems in computational biology. Single-cell RNA-seq holds great promise for powering various algorithms for network inference, but as we have shown, major obstacles remain to doing so in practice. When provided with temporally coupled measurements, Scribe accurately reconstructs networks of modest scale. As experimental and computational improvements to single-cell expression techniques couple measurements across time, we expect Scribe to be increasingly capable in dissecting the complex genetic circuits that drive development and disease.

**Code availability**. A version of Scribe (version: 0.99) used in this study is provided as Supplementary Software. The newest Scribe implemented as an R package is available through GitHub (https://github.com/cole-trapnell-lab/Scribe) CCM algorithm is implemented as the rccm package (https://github.com/cole-trapnell-lab/rccrrO which is based on https://github.com/cibavesian/rccm. The neurogenesis simulation is implemented as the scRNASeqSim package (https://github.com/cole-trapnell-lab/scRNASeaSimy Supplementary Software also includes a helper package containing helper functions as well as all analysis code that can be used to reproduce all figures and data in this study.

**Data availability**. Four public scRNA-seq data sets are used in this study. Lung dataset: GSE52583 (Treutlein et al., 2014); LPS dataset: (GSE41265); MARS-seq dataset(Paul et al., 2015): http://compgenomics.weizmann.ac.il/tanay/?pageid=649. Olsson dataset(Olsson et al., 2016): synapse id svn4975060. Live imaging dataset for the *C. elegans* is obtained from Waterston lab.

## Acknowledgements

We thank Robert Waterston and his lab for guidance in analyzing *C. elegans* early embryogenesis, Gioele La Manno for discussing causal network inference with RNA-velocity, Andysheh Mohajeri for helping preparing a website for this work, and members of the Trapnell laboratory for comments on the manuscript. This work was supported by US National Institutes of Health (NIH) grant DP2 HD088158, the Paul G. Allen Frontiers Group (Allen Discovery Center grant to CT), and the W.M. Keck Foundation (to CT). C.T. is partly supported an Alfred P. Sloan Foundation Research Fellowship.

## Contributions

X.Q., A.R., C.T. and S.K. designed Scribe. X.Q. and A.R. implemented the methods. X.Q. and A.R. performed the analysis. L. W., M. Q., T. D., and L. S. contributed to data analysis. X. Q., C.T., S. K. conceived the project. All authors wrote the manuscript.

## Supplementary Figures

**Supplementary Figure 1:**
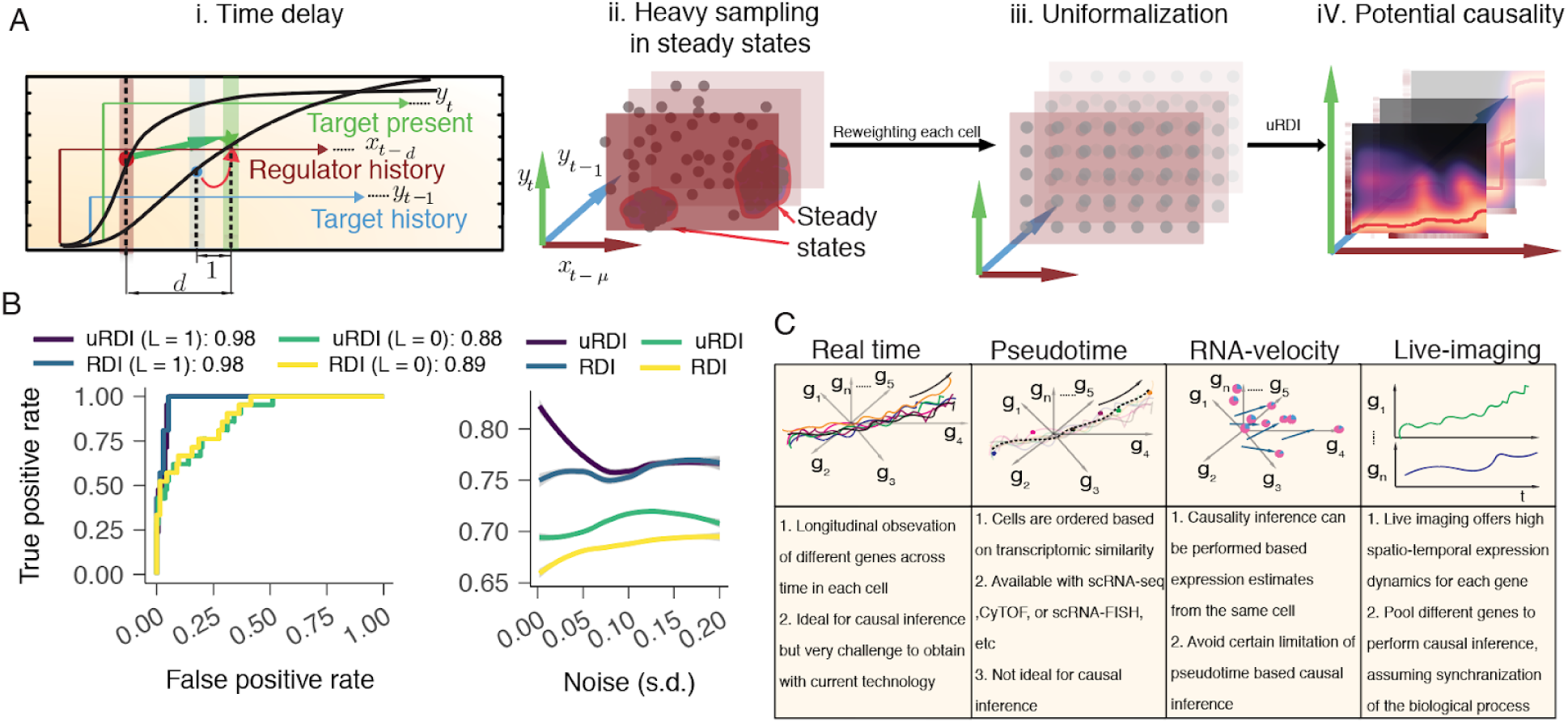
Scribe recovers causal interactions at rare transition states and generalizes to various types of single-cell time-series measurements. **(A)**. Scribe leverages a rigorous technique of uniformalization to detect potential causality. Cells often reside in the steady states but rarely in the transition state. This leads to heavier sampling of single-cell measurements in steady states (for example, the low or high expression regions at the beginning or end of the kinetic curves as shown in panel A) than transition state. More intuitively, the biased sampling in data can be seen from the state space formed with the expression values of the regulator, target and target’s history (*II*. *Heavy sampling in steady states*). In order to account for sampling biases from single-cell measures, Scribe integrates uRDI/ucRDI (uniformization of (c)RDI) by reweighting each cell and thus replacing the biased sampling distribution with a uniform distribution (*II. Uniformalization*) to rigorously quantify the *potential causality* (Rahimzamani and Kannan, 2017) (how much influence a regulator can potentially exert on target *without cognizance* to the regulator’s distribution). For a strong causal interaction, RDI requires the response of the target (see more details in **Fig 1A)** to the regulator evolves substantially under different historical states of the target (*III. Potential causality*). **(B)**. uRDI improves the recovery of causal regulations. **Left:** Receiver Operating Curves or ROC for different methods in Scribe on causal inference with or without uniformalization for the linear system. **Right:** Area Under Curve or AUC for different methods in Scribe on causal inference with or without uniformalization for the non-linear neurogenesis system. Noise for each system is treated as the same in the **Main Figure 1C**. (**C**). Scribe is generally applicable on any single-cell time-series datasets. Each column in the table corresponds to different types of time-series data. The first column corresponds to theoretically ideal real time-series datasets, where, i.e., the transcriptome for each cell is followed over time longitudinally. The second column corresponds to the pseudotime-series datasets, where the transcriptome for a population of cells at different developmental stages is captured with scRNA-seq. Using computational algorithms, for example, Monocle 2, cells are ordered to obtain pseudotime-series data. The third column corresponds to the datasets estimated from the “RNA velocity” analysis framework where the current or future mature mRNA expression, etc are estimated for each cell. In the first three columns, each axis corresponds to one gene dimension where each curve corresponds to the expression dynamics for each individual cells in the full gene space over time. The arrow points to the direction of cell differentiation and the dash line from the second figure corresponds to the inferred pseudotime trajectory. For the last column, “live-imaging” dataset, in which each cell is tracked over time but only one gene is measured, as we did for studying the *C. elegans* early embryogenesis.

**Supplementary Figure 2:**
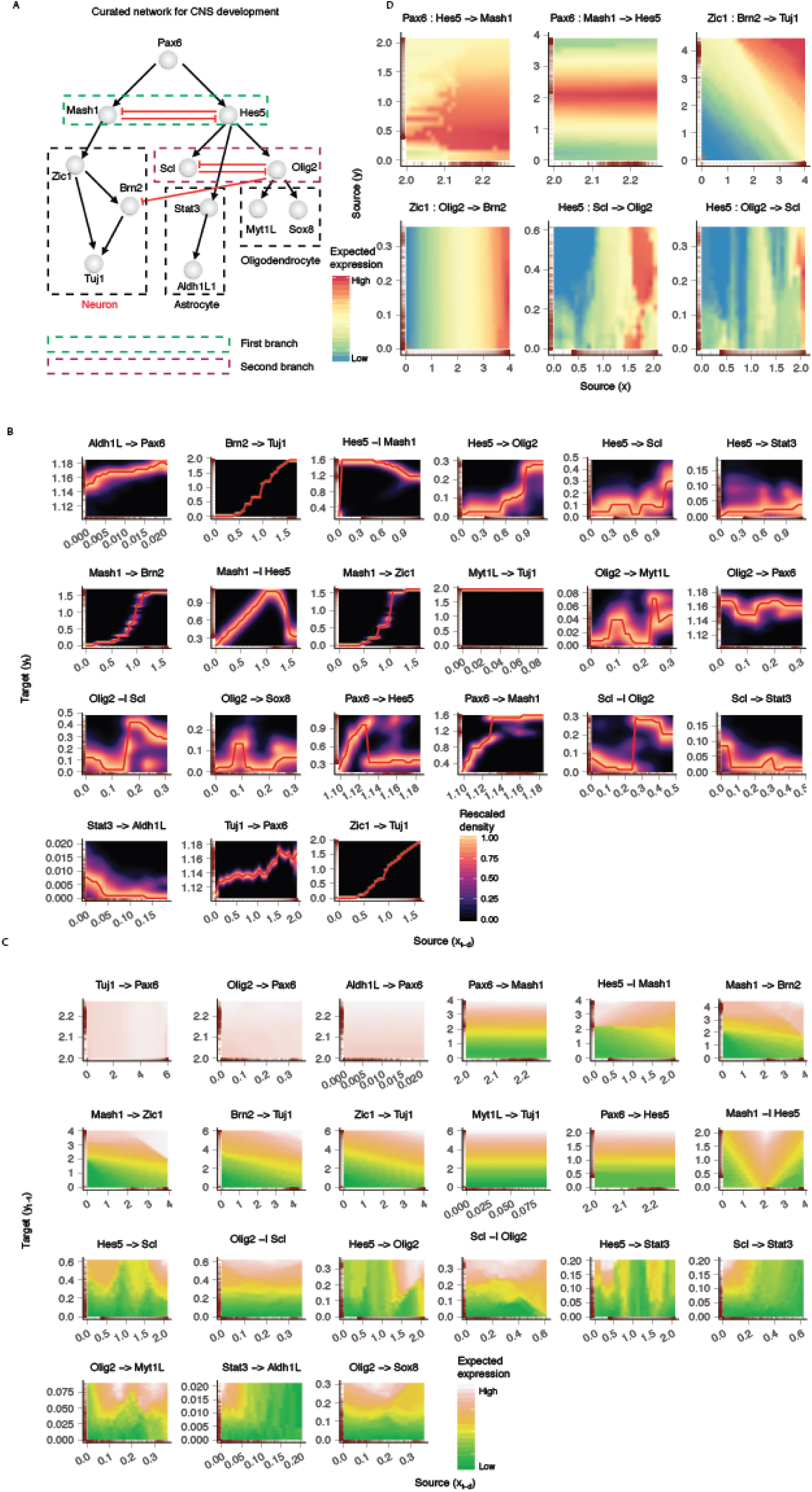
A gallery of regulatory patterns visualized by the response, causality and combinatorial regulation visualizations from Scribe on the simulated neurogenesis dataset. **(A)** The manually curated network used to simulate the three-way cell fate specification of the central nervous system (Qiu et al., 2012). The network consists of two key mutual-inhibition gene pairs. Initializing this network with small amounts of stochastic noise and following expression kinetics over time simulates the trajectory followed by a single cell leading to the fate of either neuron, astrocyte or oligodendrocyte. For simplicity, only a simulation leading to the neuron fate is used for the analysis presented in panels **B-D. (B)** Response visualization plots for all the interacting genes pairs in the network. *Response* visualization reveals the regulatory response of the target to the regulator. X-axis corresponds to the regulator’s previous expression with a time lag *d* (*x_t−d_*) while y-axis corresponds to target’s current expression (*y_t_*). For example, as shown here, response of *Brn2* to *Tuj1* is a sigmoid function suggesting positive regulation while the response of *Mash1* to *Hes5* is a threshold function suggesting threshold mutual repression. The heatmap corresponds to the rescaled normalized conditional density for two genes, similar to the DREVI plot from ref.(Krishnaswamy et al., 2014) with the difference that we explicitly consider the time-delayed response from the target to the regulator (*P* (*y_t_*|*y_t−d_*)). The red line represents the most probable value for the target given a regulator’s expression. The rug plot on the axis corresponds to the density of cells at a particular value on the corresponding dimension. **(C)** Causality visualization plots for all interacting genes pairs in the network. **(D)** Combinatorial logic visualization plots for all six two-input combinatorial regulations cases in the network.

**Supplementary Figure 3:**
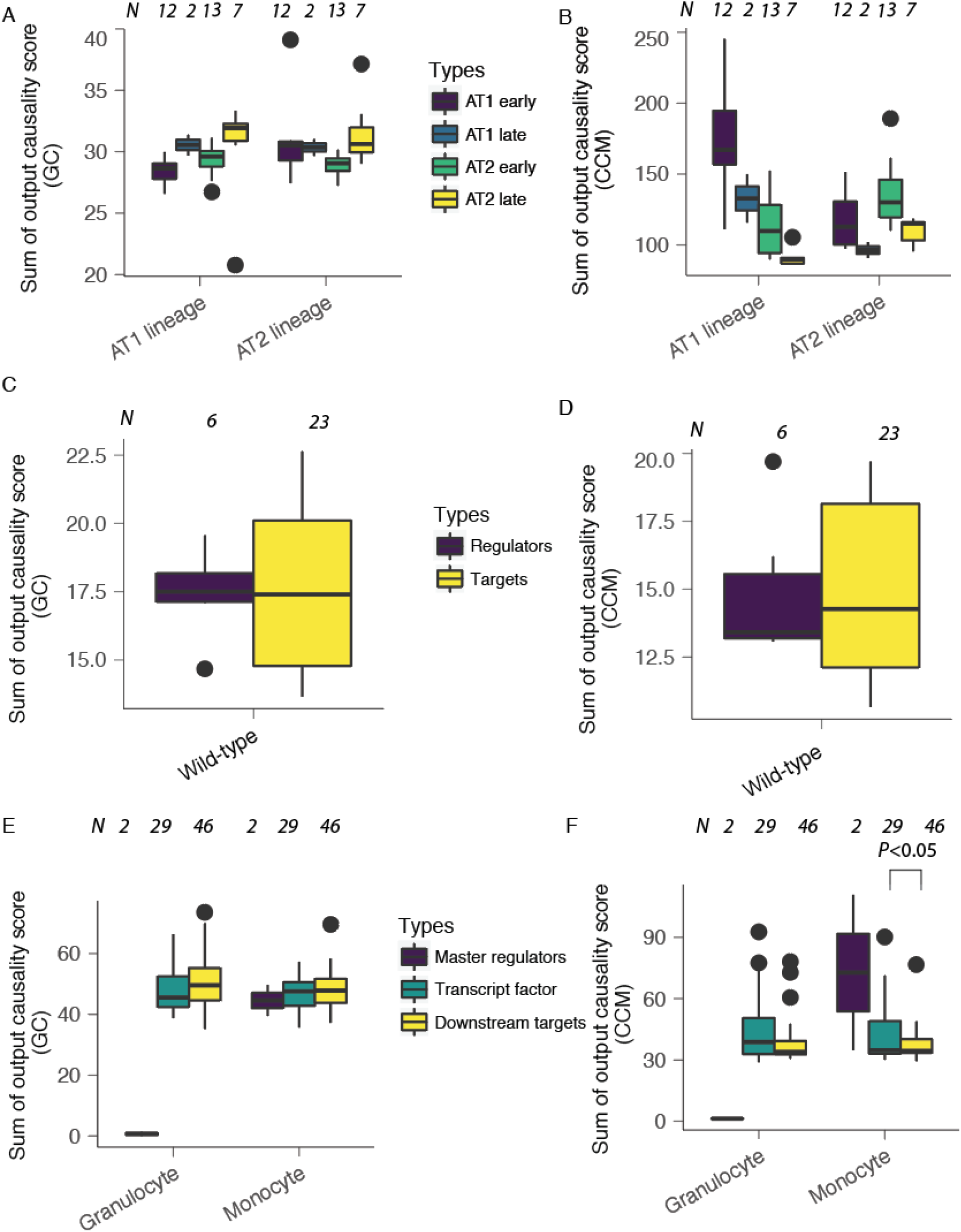
Poor performance of other causal inference algorithms in resolving gene regulatory directionality. To test whether or not other well-known causality detection algorithms (Granger Causality (GC) and Convergent Cross Mapping (CCM)) can also infer the correct regulatory directionality, we calculated the total outgoing causality scores inferred from them for the known regulators and targets on the same datasets as used in main text **Fig 5. (A, B)** Distribution of total outgoing causality scores compared to that of the target genes across AT1 or AT2 branch based on GC **(A, left)** or CCM **(B, right). (C, D)** the same as in **(A, B)** but for the LPS data (wild-type cell subset). **(E, F)** The same as in **(A, B)** but across granulocyte and monocyte branches of the Olsson dataset (wild-type cell subset). To calculate total causality score, causal strength between all the genes are calculated with RDI which is then normalized with CLR algorithm.

**Supplementary Figure 4:**
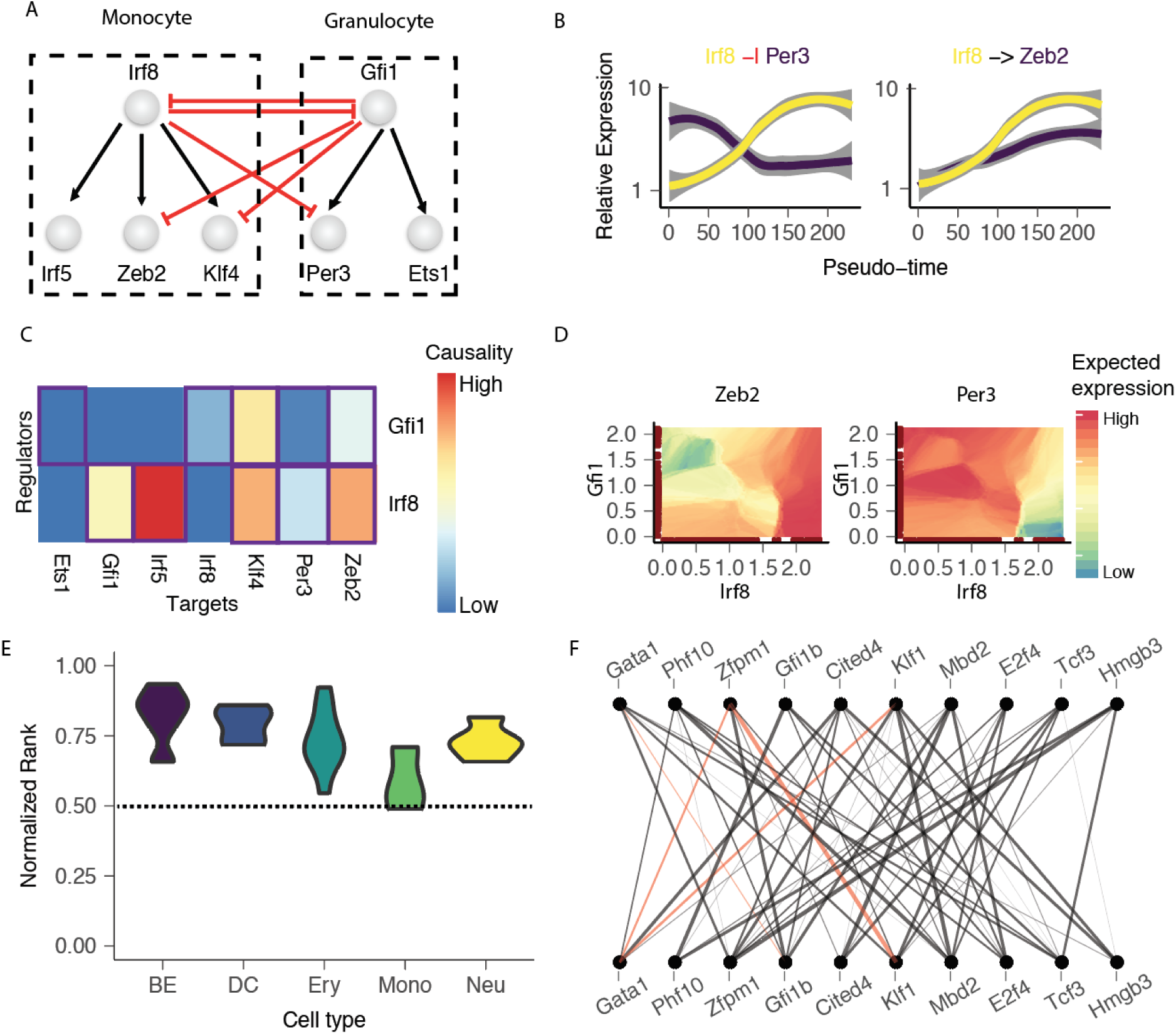
Scribe recovers a core regulatory network responsible for myelopoiesis. **(A)** A core network describes key regulators during the specification of monocytes and granulocytes based on data collected from perturbation experiments, bulk ATAC-seq and ChIP-seq data(Olsson et al., 2016). **(B)** Examples of gene-target pair kinetic curves over pseudotime along the monocyte lineage. **(C)** Scribe infers the expected core regulatory network interactions for myelopoiesis. Causal scores from regulators to all other genes are calculated using RDI and are then normalized using the CLR algorithm. **(D)** Visualization of combinatorial gene regulation from *Irf8* and *Gfi1* to *Zeb2* or *Pert*. Gene expression values are denoised through reversed graph embedding (Qiu et al., 2017a) and calculated as a local average. Values are then rasterized to plot as a two-dimensional heatmap (See **methods** for details). **(E)** Normalized rank of lineage specific genes’ total outgoing RDI sum. Total outgoing RDI sum is calculated for all lineage specific genes for each lineage as in **Figure 5**. The normalized rank is calculated based on the order of each lineage specific TF among all significant branching TFs divided by its total number. When the normalized rank is close to 1, the corresponding gene is close to have the highest sum of outgoing RDI scores. The dash line indicates the average rank (0.5) for a random gene. **(F)** Lineage specific network of significant regulators during erythropoiesis. Edges supported by SPRING database is colored as red lines. For panels **E (F)**, BEAM analysis was used to identify significant branching genes associated with the four (one) lineage bifurcation events shown in the haematopoietic trajectory from ref. (Qiu et al., 2017a) based on the paul dataset (Paul et al., 2015). The top 1,000 differentially expressed genes associated with each bifurcation were chosen to build a causal network for each relevant lineage. A set of TFs relevant to specific lineages described previously are used for panel **E** or **F**. Neu: Neutrophil; Ery: Erythroid, Mk: Megakaryocyte; Mono: Monocyte; DC: Dendritic Cell; BE: Basophil / Eosinophil.

**Supplementary Figure 5:**
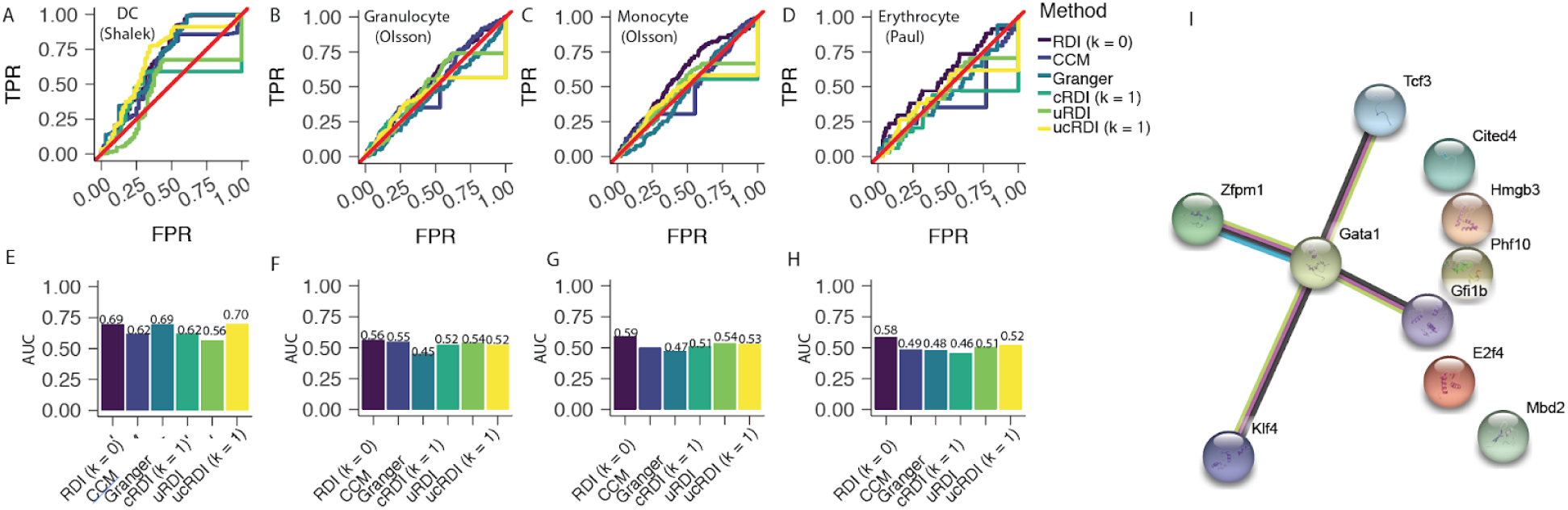
Benchmark Scribe with other leading causality detection algorithms. **(A, E)** Receiver Operating Curves or ROC **(A, top)** and Area Under Curve or AUC **(E, bottom)** of the inferred causal network based on Scribe, GC and CCM on the Dendritic Cells (DC) dataset. Four different variants of causal inference implemented in Scribe are tested: *RDI* (*L = 0*): the default RDI method without conditioning on any other gene; *RDI* (*L = 1*): the RDI method based on conditioning on the incoming gene with highest causality score, except the current target; *uRDI:* the method based on the uniformization technique applied on the actual distribution in RDI; *uRDI* (*L = 1*): the uRDI method but also with the conditioning on the incoming gene with the highest causality score, except the current target. The network from(Amit et al., 2009) based on a unbiased perturbation experiment is used as benchmark gold-standard. **(B, F)** The same as in **(A, E)** but for the granulocyte branch of the Olsson dataset. **(C, G)** The same as in **(A, E)** but for the monocyte branch of the Olsson dataset. The manually curated network for the myeloid differentiation from (Su et al., 2017) is used as the benchmark gold-standard. **(D, H)** The same as in **(A, E)** but for the erythroid branch of the Paul dataset. All wild-type cells are pooled to reconstruct the developmental trajectory and a subset of CMP and erythroid branch cells are used to estimate the causal network for the erythroid branch. The manually curated network for the erythroid differentiation from Ref. (Swiers et al., 2006) is used as benchmark gold-standard. **(I)** The network of the gene-set as included in the panel **(Supplementary Figure 4F)** retrieved from the STRING database. See https://strina-db.org/cai/network.pl?taskld=2QGoh9uvYldY for more details.

**Supplementary Figure 6:**
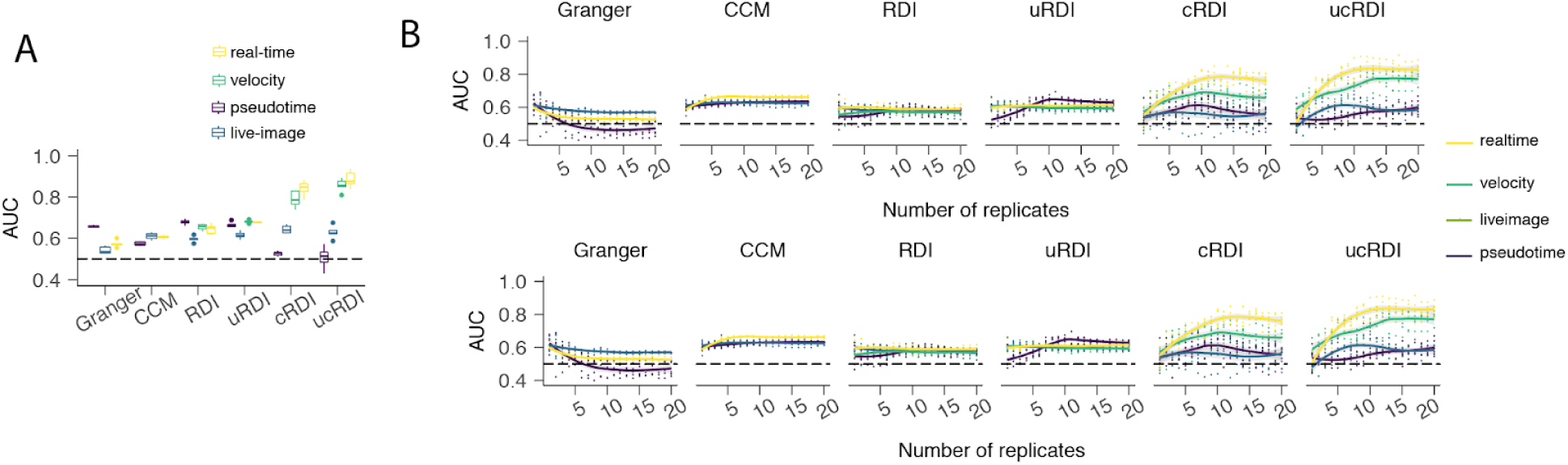
Benchmark different causal inference methods with different types of single-cell time-series datasets and under different number of replicates of developmental trajectories. **(A)** Dynamics coupling in real time and “RNA velocity” datasets enables Scribe to correctly infer causal regulatory network. Scribe (including four variants, RDI ( *k* = 0), uRDI (*k = 0*), RDI (*k =* 1) and uRDI (*k =* 1)) and two alternative causal inference methods, Granger and CCM are used to reconstruct network based on 2,000 data points from each of the four different single-cell time-series datasets. The results from each method are then compared with the known network architecture to obtain the AUC score. The data generated with time delay between any regulator to any target ranges from 1 to 3. **(B)** The same analysis as in main text **Fig 6C** is performed but downsampling datasets in terms of the number of replicates of developmental trajectories (see **Methods** for more details). The top panel corresponds to the data generated with time delay between any regulator to any target as 1 while the bottom panel with time delay for different regulators ranging from 1 to 3.

**Supplementary Figure 7:**
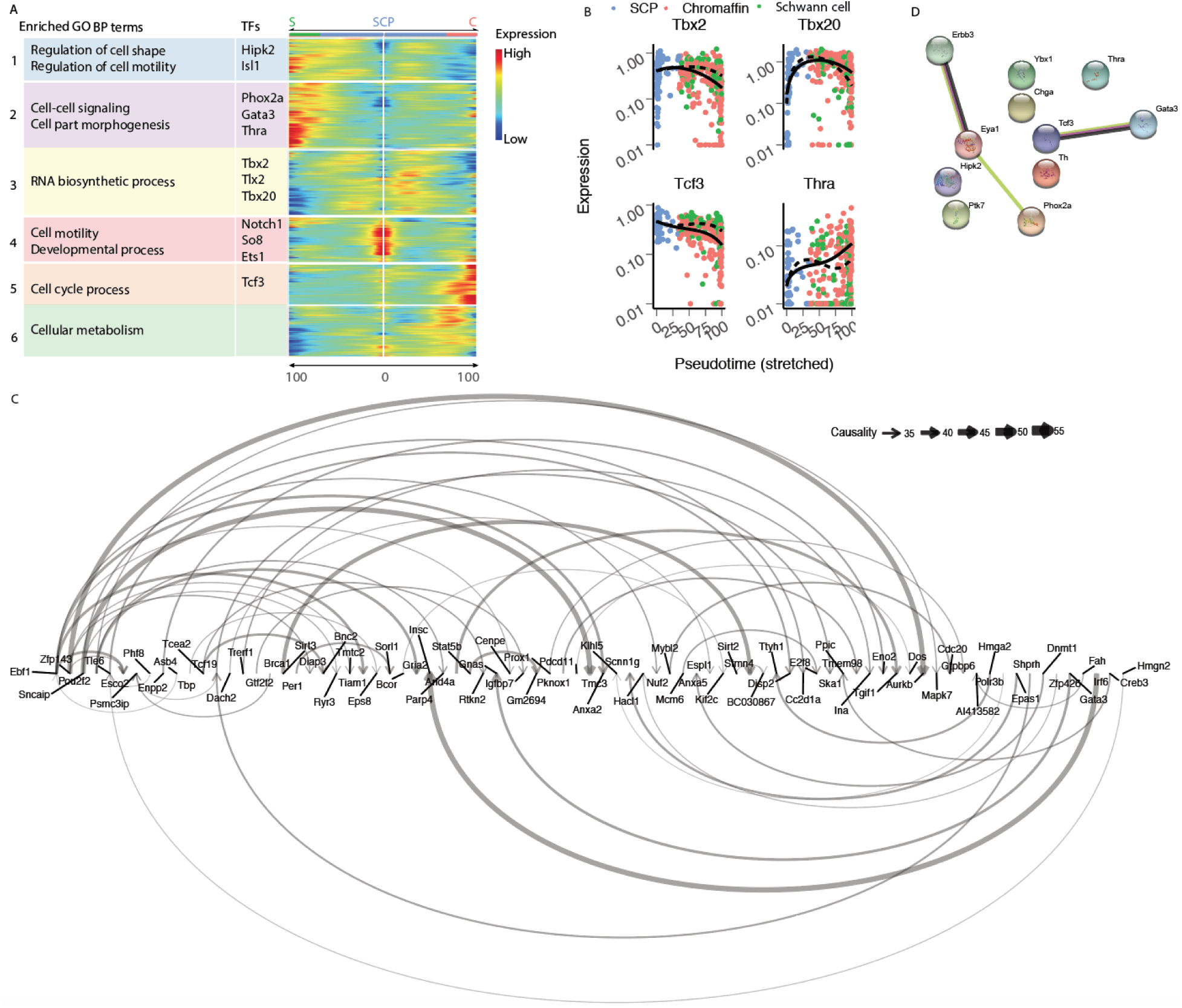
Comparing causal inference based on pseudotime or RNA-velocity. **(A)** Clusters of significant branching genes reveal distinct transcriptional programs between Schwann and chromaffin cell lineages. **(B)** Gene expression dynamics of representative significant branching transcription factors from **(A). (C)** Top 100 causal edges between significant branching transcription factors (TFs) as well as from TFs to the potential targets in chromaffin lineage inferred with RNA-velocity. **(D)** The network of the gene-set as included in the **Fig 7C** retrieved from the STRING database. For more details, please see https://string-db.org/cgi/network.pl?taskId=d3PMC9KwRuep.

## Methods

### Four possible single-cell time-series measurement modalities

Cell differentiation is an intrinsically noisy and asynchronous process. Even for the same developmental process, every cell in any given time should be regarded as a distinct sample.

We consider four possible types of gene expression measurements in those single-cell samples:

1. Real-time, where we measure the gene expression for all genes simultaneously in a single cell over time. This is the ideal situation but no existing technology can produce data like this yet.
2. “RNA-velocity” where we only capture the current state and the next state for all genes in different cells. “RNA-velocity” can be computationally inferred from single-cell RNA-seq datasets, or directly measured with Seq-FISH(Shah et al., 2018), and possibly single-cell version of SLAM-seq (Herzog et al., 2017; Muhar et al., 2018), TUC-seq (Riml et al., 2017) and TimeLapse-seq (Schofield et al., 2018), among others.
3. Live-imaging datasets are those generated with multiple separate live-imagings for a single protein in a single-cell which are then aligned along the same developmental process to form a time-series for all genes.
4. Pseudotime is where we apply trajectory reconstruction algorithm to order the single-cell RNA-seq snapshot dataset to form a time-series.

### The problem of causal regulatory network inference

In this work, we formulate the problem of causal regulatory network inference as the inference of the underlying structure of influences in a stochastic dynamical system where the time series of each gene is causally regulated by a subset of other genes. We assume that there are no unobserved confounders in order to make the problem tractable. In this setting, we can potentially infer the causal regulators based on estimating the amount of information transferred from one variable (a potential regulator) to another time-delayed response variable (a potential target). In the context of single cell genomics (e.g. scRNA-seq, live cell imaging), we ask how we can reconstruct a regulatory network consisting of causal regulations that accurately describe the gene expression dynamics and the associated cell fate transitions.

### Causal Inference

In the setting stated above, various techniques, including Granger Causality and CCM, each associated with different assumptions have been proposed to detect the structure of the causal regulatory network. In the following, we briefly summarize these methods and introduce RDI, the method we developed and used in this study.

#### Granger causality

In order to determine whether one time series (*X*_1_) is useful in forecasting another (*X*_2_) in economics, Clive Granger first proposed Granger Causality (GC) in 1969 (Granger, 1969). According to GC, if *X*_1_ "Granger causes" *X*_2_, then the predictability of *X*_2_ based on past values of *X*_2_ and *X*_1_ together is significantly greater than that of predicting purely based on the past values of *X*_2_. GC in its original formulation, however, is only able to detect linear causal regulation: i.e., when the regulators regulate the target through a linear relationship.

### Convergent Cross Mapping

In order to detect pairwise non-linear interactions in deterministic ecology systems, George Sugihara and colleagues proposed Convergent Cross Mapping (CCM) which is based on state-space reconstruction (Sugihara et al., 2012). One fundamental and somewhat counterintuitive idea of CCM, distinct from GC, is that it is possible to estimate *X*_1_ from *X*_2_, but not the other way if causation is from *X*_1_ to *X*_1_. CCM first constructs shadow manifolds 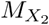 and 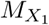 from lagged coordinates of the time-series *X*_2_ and *X*_1_. It then tests whether states in the shadow manifold 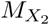 can be used for estimating the states in 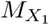 and *vice versa* via mapping through nearest neighbors (*cross mapping*). Another key idea of CCM is *convergence* which means that as the length of the time-series increases, the shadow manifolds become denser and the ellipsoid or space formed by nearest neighbors shrinks, leading to improvement of cross-map estimates. Although CCM is appealing, it cannot be generalized to stochastic systems as Takens’ theorem, the cornerstone of CCM, will break down in such scenarios (Takens, 1981). Furthermore, CCM can only infer pairwise relationships and complex multi-factorial interactions common in gene regulatory networks are not captured in CCM.

### Restricted Directed Information (RDI)

As mentioned earlier, the causal inference method in Scribe is based on Restricted Directed Information (RDI). This measure determines the amount of *statistical inter-dependence* (or more formally the *mutual information*) between the past state of the regulator and current state of the target gene conditioned on the target’s immediate previous state.

Cell state transitions are controlled by hierarchical regulatory networks (Peter and Davidson, 2011). In such networks, as the expression of regulator changes, their downstream target responds accordingly after some time delay *d*. A canonical measure of mutual dependence which accounts for both linear and nonlinear associations between two genes (or more generally, two random variables)*, X, Y*, is mutual information (MI)(Cover, 2006). MI is symmetric and can quantify the "amount of information" obtained about gene *X* or *Y*, through the other gene *Y* or *X*. It essentially determines how similar the joint distribution (*PXY*(*x,y*)) of the two genes *X, Y* is to the products of factored marginal distribution *pX*(*x*)*pY*(*y*), or formally:

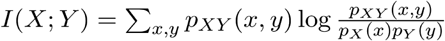

If *I*(*X;Y*) is zero, then the two genes *X, Y* are independent; otherwise it implies there exists some dependency between them (e.g. in the case of a regulator and its target). It is often useful to quantify the mutual dependence between two random variables (for example, regulator *X* and target *Y*) while removing the effect of a third random variable (for example another regulator *Z* or the history state of the target). This leads to developing of conditional mutual information, which is defined as:

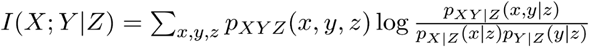

Ml provides a powerful approach to quantify the symmetric interdependence between genes. However, a favorable approach would be to measure the causal score from a potential regulator to its target. We can achieve this by considering the time-series of regulators and targets 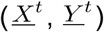 and quantifying the information transfer from the past state(s) of *X* to the current state of the variable *Y* denoted by *Y*(*t*).

Previously, T. Schreiber reported *Directed Information* (*DI*) as a measure for the amount of information flowing from the past state(s) of *X*, the regulator, to the the current state of the variable *Y*, the target (Schreiber, 2000). DI is defined as:

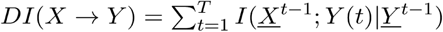

In order to remove indirect interactions, we can calculate the information transferred from the regulator to the target while conditioning on all the other genes ({*X_i_, X_j_*}*^c^*), which is,

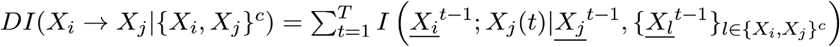

Furthermore, for a set of genes of interest, *X*_1_*,X*_2_*,…,X_N_*, from a single-cell genomics dataset, we can infer a Directed Information graph, *G_DI_ =* (*V, E*) where the vertex set *V* corresponds to the genes {*X*_1_*,X*_2_*,…*, *X_N_*} and the edge *e_ij_* = (*X_i_*, *X_j_*) from gene *X_i_* to *X_j_* exists if and only if *DI*(*X_i_* →· *X_j_*|{*X_i_, X_j_*}*^c^*) ≠ 0 the edge weight corresponds to the quantified DI value *DI*(*X_i_* →· *X_j_*|{*X_i_, X_j_*}*^c^*).

It was shown that if a system is not purely deterministic, the directed information graph *G_DI_* inferred from DI will correctly recover the true causal graph *G_C_* (the network which includes all causal interactions as directed edges) (Sun et al., 2015). Although DI is able to detect both linear and non-linear causality as opposed to the linear Granger causality and is applicable to stochastic systems, it (1) can not deal with deterministic systems which may be of interest for certain scenarios and (2) poses huge computational burden because it conditions on all possible previous states of the regulator or target and (3) requires enormous amount of data which is not affordable even with current single-cell genomic datasets.

We recently proposed a novel formulation of DI to alleviate those issues by employing only the immediate past of the target or regulators instead of all the past states assuming a first-order Markov system, which is generally applicable to most biological processes. In this method, the randomness is present due to the random initialization of the Markov system, hence creating a random process on which information measures are well defined. We term this novel method as Restricted Directed Information (RDI) and define it as,

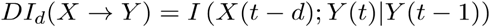

Despite the fact that original RDI measure is defined only for the immediate past of the regulator *X*, this measure can be flexibly defined for arbitrary effect delay *d* from *X* to *Y* as we have done here.

*Conditional Restricted Directed Information* (*cRDI*): Similar to (Schreiber, 2000), RDI can also be extended to the case where the information transfer from *X* to *Y* is conditioned on other potential regulator(s) *Z* to rule out the possible indirect causal effects and confounding factors. Thus the Conditional RDI (abbreviated as cRDI) can be formulated as:

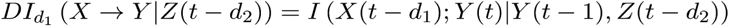

In (Rahimzamani, *et. al*, Allerton 2016), it’s shown that cRDI works in many stochastic or deterministic cases and under some mild assumptions is capable of inferring the correct regulatory network *G_C_*. Moreover, it has shown that if the conditions are violated, no other method will be able to recover the correct network (see Section IV. in (Rahimzamani, *et. al*, Allerton 2016)).

In the upcoming sections we will discuss how RDI and cRDI are utilized in the Scribe toolkit.

### Uniformization method for adjusting sampling bias

During our studies over the simulated benchmark data, we found that as the number of samples increases, performance of RDI first increases and then start to decrease. This problem was particularly acute in simulations where gene expression reached a plateau after cells committing to a cell fate. In general, while the transitional states are of higher importance in discovery of causal interactions, oversampled equilibrium states will outnumber the transitional samples resulting in a sampling bias towards less informative equilibrium states. This phenomenon can in turn reduce the inference accuracy, since RDI requires calculating conditional mutual information (*I*(*X*(*t − d*)*;Y*(*t*)|*Y*(*t* − 1))) by design, which is a function of the joint distribution (*p*(*x_t−d_,y_t_,y_t−_*_1_) *=p*(*y_t_*|*x_t−d_, y_t−_*_1_)*p*(*x_t−d_,y_t−_*_1_)). That is, the distribution is influential in the RDI calculation, despite the fact that the RDI score should be fully determined only by the conditional distribution. Hence we devised a scheme to correct for sampling bias by re-weighting samples so that those from the system during transitional periods are weighted higher than cells sampled from the system at equilibrium. One may assume the input distribution is uniform and redistribute the observed samples in a more homogeneous fashion before calculating the RDI value.

This bias correction scheme, which we term *Uniformized conditional mutual information* (uCMI) replaces the actual distribution *p*(*x_t−d_*, *y_t−_*_1_) with a uniform distribution *u*(*x_t−d_,y_t−_*_1_) and then calculates the conditional mutual information for *p*(*y_t_*|*x_t-d_,y_t−_*_1_)*u*(*x_t-d_,y_t−_*_1_). This is made possible thanks to the concept of *potential Conditional Mutual Information* (qCMI) (Rahimzamani and Kannan, 2017) and a novel estimator, in which the actual distribution *p*(*x_t-d_, y_t−_*_1_) of samples is replaced by any arbitrary distribution *q*(*x_t−d_, y_t−_*_1_) before estimating the conditional mutual information. uCMI is thus a special case of qCMI, in which the replacement distribution *q*(*x_t-d_,y_t−_*_1_) is uniform. By replacing the conditional mutual information (CMI) in RDI with uCMI, we obtain a new way of computing information transfer called *uniformized Restricted Directed Information* (uRDI).

The discussion above is especially relevant for single cell genomics datasets as single cells are not homogeneously spread across many biological processes and they often will be heavily sampled from steady states while rarely from transition states. A compelling discussion of this phenomenon can be found in c.f. (Olsson et al., 2016). This imbalance of sampling confounds the performance of RDI (or other mutual information based methods) and thus leads to ignorance of rare but critical regulation happened during transition states. We noticed that empirical methods have been reported to account for sampling biases from single-cell measures (Krishnaswamy et al., 2014). However, the uRDI method incorporated in Scribe provides a rigorous approach to replace the biased sampling distribution with a uniform distribution to quantify potential causality (how much influence a regulator can potentially exert on target without cognizance to the regulator’s distribution) and is thus arguably a superior approach to account for the sampling biases issue (Rahimzamani and Kannan, 2017).

### Scribe: a toolkit for visualization and detection of complex causal regulation from single cell genomics datasets

Although Scribe is applicable to any time-series datasets, it is specifically designed for visualizing and detecting complex gene regulation from single cell genomics datasets (e.g. scRNA-seq). Scribe relies on (uniformized) restricted directed information to detect causality but also supports other methods, including the well-known mutual information, Granger causality and the more recent CCM. Scribe starts with time-series data, which can be based on “pseudotime-series”of a developmental trajectory reconstructed from scRNA-seq data such as those constructed using Monocle 2, live imaging data or datasets with current and predicted spliced RNA expression estimated using RNA-velocity. Scribe provides two main types of analysis:

1. Visualization and estimation of causal gene regulation;
2. Reconstruction of large-scale sparse causal regulatory networks.

#### Preparing pseudotime-series or RNA-velocity for scRNA-seq datasets

Scribe does not provide any built-in functionalities for pseudotime-series construction and relies on Monocle (http://cole-trapnell-lab.github.io/monocle-release/) or similar tools, such as dpt(Haghverdi et al., 2016) or wishbone(Setty et al., 2016), for reconstructing the single-cell trajectory before inferring causal networks. Scribe also doesn’t provide any built-in functionalities for RNA-velocity estimation and relies on the velocyto framework (La Manno et al., 2017) for those estimations. In relation to physical time, pseudotime has an arbitrary scale, thus Scribe doesn't consider pseudotime value themselves instead using the ordering of each cell in pseudotime for causal network inference. Similarly, we also assuming the time delays Δ*t* used in RNA-velocity estimations are constant across cell and genes for the sake of simplicity.

#### Visualizing pairwise gene interaction

In order to intuitively visualize casual regulations between genes, Scribe provides different strategies to visualize the **response, causality** and **combinatorial regulatory logic between gene pairs**. The response visualization is similar to the DREVI approach as proposed by Smita Krishnaswamy, et. al(Krishnaswamy et al., 2014) with the exception that it considers time delay to visualize the expected expression of potential targets given a potential regulator’s expression after a time delay. Response visualization thus additionally aids in visualizing commonly appeared time-delayed regulations involved in cell differentiation(Alon, 2007).

One limitation of response visualization is that it ignores the effects of a gene's previous state to the current state or memory of its history. In order to also capture this effect and thus intuitively visualize causality, Scribe is equipped with causality visualization. Essentially, this approach visualizes the causal regulation by considering the information transfer from the time delayed potential regulator to the target’s current expression, conditioned on the target’s previous state to remove effects from auto-regulation. Causality visualization is a heatmap consisting of the expected value of the target's current expression given the target’s immediate past expression (y-axis) and regulator’s expression with a time lag *d* (x-axis). For each column, it represents the relationship for the target's expression at the previous time point to the current state (memory of the history or “auto-regulation”) given a fixed regulator value, while for each row, the information transfer from the regulator to its targets given the previous target state.

#### Visualizing combinatorial gene regulation

It is of great interest to understand the combinatorial gene regulation as it often determines how cells make decisions to choose a particular cell fate or adapt to external stimuli(Ma et al., 2009). In order to visualize two-input combinatorial regulation, Scribe provides a third visualization tool. This visualization is a heatmap consisting of the expected value of the target's current expression given knowledge of both of the regulators' expressions with a time lag (x/y-axis). For both of the causality and the combinatorial logic visualizations, the corresponding expected value is calculated through a local average with a Gaussian kernel.

We noticed that gene regulation *directly* affects the rate of the target gene which then results in gene expression changes. For example, if a gene *x* is negatively regulated by gene *y*. We may define the rate function of *x*: as 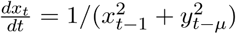. Therefore, visualizing the expected rate of a target at its current state given knowledge of both the regulators' expression with a time lag (x/y-axis) allows better intuition of regulations. Although we won’t have accurate estimates of the rate of gene expression with pseudotime-series data, the RNA-velocity method can be used to obtain those estimates.

We also noted that combinatorial gene regulation visualization is especially useful to help us visually identify potential direct/indirect regulators. For example, if we have a regulatory pathway *x− > y− > z*, we will see that the expected gene expression of *z* is only dependent on the direct regulator *y* instead of the indirect regulator *x* from the visualization **(Figure 2D)**. That is, combinatorial gene regulation visualization indeed provides visual intuitions for the conditional RDI (in the above case, the row corresponds to *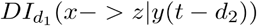* while the column 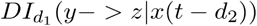)

#### Causal network inference: an RDI-based algorithm

Causal inference in Scribe is based on RDI, which is an extension of directed information under the assumption that the underlying processes can be described by a first order Markov model. The method we implemented basically tries to calculate the RDI value for each pair of genes (*i, j*) conditioned over the top *L* genes (default is 0 or no conditioning and 1 for cases where we used conditioning) which are candidates of being regulators of the gene *j*.

To reach this goal, it first calculates all the pairwise *unconditioned* RDI values, for all the potential delays specified by the user in vector *d* (by default, it is a vector including 5, 10, 20, 25). Note that for RNA-velocity dataset, since we assume the time delays Δ*t* for the current and predicted future RNA expression level are constant across the cell and genes, there is no need to scan for a window of potential time delays. Then for each pair it treats the delay corresponding to the largest RDI value as the *“true” delay of effect*, i.e. the actual time delay by which the effect of *i* appears in *i*. Having identified the “true” delays, the method then re-calculates the pairwise RDI values for each pair of genes (*i, j*), this time conditioned over the top *L* (*L* can be specified by the user) genes with the highest incoming RDI values to *j* associated with their corresponding true delays, treating them as the potential regulators of *j*.

The algorithm of causal inference in Scribe is as follows:

**Input:** gene expression time-series (either based on pseudotime-series, “RNA-velocity” or living imaging data, among others) *X_i_* for each gene *i*

**Output:** A matrix of pairwise causality scores

**Parameters:** *d:* vector of delays, *L*: number of conditioning genes

**Pseudocode:**

1. For each pair of genes (*i, j*)

- For all delays *δ ∈ d*: Calculate *RDI_δ_*(*X_i_* → *X_j_*)
- Set 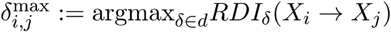
2. For each gene *j*:

- For all *i*: sort 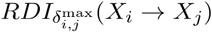 values in descending order
- According to the sorting above, take the *L +* 1 nodes *i* with the highest incoming RDI values to *j* and store them in a set as 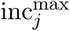. Store their corresponding delays 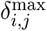 in a set 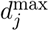.
3. For each pair of genes (*i, j*)

- If 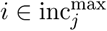 remove *i* from 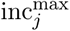. Otherwise, remove the node with the lowest 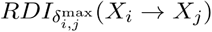from 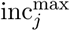
4. output 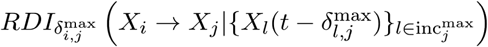

The estimation of the mutual information is inspired by Kraskov’s method (Kraskov et al., 2004), which builds on counting nearest-neighbor points. In the R implementation of Scribe, nearest-neighbor points are identified with a modified RANN package.

To calculate the causal network with uRDI, we apply the same algorithm as above but simply replace RDI with uRDI. In addition to what required in RDI, uRDI also needs to estimate the actual distribution, *p*(*x_t-d_, y_t−_*_1_), which relies on kernel density estimation (KDE). We use standard Gaussian kernels from R in the Scribe package to calculate KDE.

#### Inferring and visualizing transcriptomic gene regulatory network

Scribe can estimate a causal network from a set of known TFs (and among the TFs) to a set of targets of interest (selected through, for example the BEAM test), or estimate the pairwise causality among all the genes in a set of genes of interest. For the first scenario, Scribe estimates causality between all pairs of TFs, and the causality from each TF to each putative target; for the second scenario, Scribe estimates causality for any pair of genes in both directions. In order to retrieve significant causal edges while removing promiscuous edges and reconstruct a sparse causal regulatory network that satisfies known properties of biology networks, Scribe relies on a modified CLR method (*Context Likelihood of Relatedness*) and a novel directed network regularization inspired by some biological assumptions (see section ***Network sparsifier: CLR and directed graph regularization*** below).

In order to facilitate the visualization of complex networks, Scribe provides a variety of approaches to visualize the RDI network either through a heatmap, a hierarchical layout, an arc diagram or a hive plot, implemented based on *igraph*, *netbiov*, *ggraph, arcdiagram* as well as the *HiveR* R packages.

We used the Kleinberg centrality to define the hubness used to order genes on the arc plot which is defined as the the principal eigenvector of *A* * *t*(*A*), where *A* is the adjacency matrix of the graph(Kleinberg, 1999).

In addition to the core causality detection feature based on (uniformalized) restricted direction information, Scribe also supports various methods for inferring the regulatory relationships including mutual information, Granger causality and CCM implemented based on *parmigene, vars* and the *rEDM* packages, respectively. We also provide a python package for most of estimation methods, although without extensive support for visualization which may be supported in future.

### Parameters of RDI

| Parameter | Type | Effect of tuning parameters |
| --- | --- | --- |
| **d** | **Vector of positive integers** | Default: 5, 20, 40<br>The vector of potential delays, for which the corresponding RDI values are calculated.<br>Setting this argument too small may limit the ability of Scribe to detect causal relationships, while setting it too large can result in discovery of incorrect or indirect causal relationships, resulting in false delays and conditioning. |
| **L** | **Non-negative Integer** | Default: 0<br>Number of the top incoming node(s) to the target, excluding the source, over which RDI is conditioned.<br>L=0 corresponds to no conditioning (Plain pair-wise RDI). Any L>0 corresponds to conditional RDI (cRDI).<br>Conditioning over more nodes approaches the theoretical prerequisite of conditioning over all genes, excluding the source and target, needed for inferring the true causal network, however it imposes more computational burden and undesirably reduces the accuracy of the RDI estimator with fixed number of samples N, as it exponentially increases the dimension of the state space used to calculate the k-nearest neighbors. |
| **k** | **Positive Integer** | Default: 5<br>Number of the nearest neighbors in kNN estimator for the conditional mutual information. The parameter should be set in such a way so the neighborhood captures adequate number of samples for a good estimate of the probability corresponding to each sample. |
| **Uniformization** | **Boolean** | Default: False<br>If True, uRDI instead of RDI will be used. While imposing higher computational burden over the same data than RDI, uRDI is expected to improve the causal inference in the cases with highly-biased sampling distributions. |

### Algorithm complexity

| Algorithm | Methodology | Parameters | Complexity<br>N: the number of samples;<br>d: the dimension of the manifolds (default 2);<br>k: the number of nearest neighbors<br>L: number of<br>I: the dimension of the features data |
| --- | --- | --- | --- |
| **CCM** | Determining the causality from X to Y based on how well one can reconstruct the cross-mapped estimate of X from the nearest neighbors determined on Y space | **E**: The number of lags embedded in the shadow manifold<br>**Tau**: The time lag between each consecutive pair of time samples (default: 1) | $O(2Nd+2kN)+O(N)$ |
| **Granger Causality** | Determining the causality from X to Y based on how much the past samples of X contribute in linearly estimating the current state of Y, compared to when the Y is estimated based merely upon its own past | **Maxlag**: The number of lags of past sample included in estimating the current state of Y | $O(I^2 N + I^3 + I^2 N + I N)$ |
| **RDI and cRDI** | Determining the causality from X to Y based on the amount of mutual information between the past of X and the current state of Y conditioned over the past of (potentially) all other variables than X | **k**: The number of neighbors for kNN estimation of mutual information<br>**Delays**: The lags for which the mutual information from the lagged source to the current state of target is estimated.<br>**L**: The number of the conditioning nodes other than X and Y | $O(dN \log N + kN) + O(N)$ |
| **uRDI and ucRDI** | Same as RDI method, but including the replacement of the empirical distribution of the past samples with a uniform distribution | **All Parameters from RDI plus:**<br>**BW:** The bandwidth of the kernel estimator | $O(dN \log N + kN) + O(N^3) + O(N)$ |

### Regularizing causal interaction networks

In theory, Scribe can remove potential indirect causal gene regulation from one gene *x* to another gene *y* by conditioning on all other genes in the transcriptome except *x*. However, this requires huge amount of samples which is infeasible even with current single cell genomics techniques and is impractically slow for even modest sets of genes. Therefore, we sought alternative approaches based on statistical significance and reasonable assumptions of biology structures to remove potential indirect edges. The first method we applied is the CLR or *Context Likelihood Relatedness*. After computing the causality score with RDI (uRDI) without conditioning between all gene-pairs, CLR calculates a normalized score based on the z-score (or 0 if the z-score is less than 0) from all the input edges to the potential target and all the output edges from the potential regulator of the gene pair. This normalized score is used as a statistical likelihood of each causal edge regarding to its network context. More formally, denoting the asymmetric matrix *R* corresponds to all raw causality scores calculated with Scribe, with *R_ij_* being the causality score from gene *i* to gene *j*, we can calculate the z-score *z_i_* based on all gene *i*’s output causality scores and *z_j_* all gene *j*’s input causality scores. The normalized score of *R_ij_*, 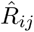 is defined as:

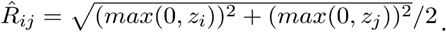

The user can either use the normalized score or choose a threshold of the normalized scores and treat the edges above the threshold as significant or real regulation comparing to the background distribution of the causality scores. As discussed in the original study, CLR removes many of the false regulations in the network by eliminating “promiscuous” cases, where one regulator weakly co-varies with a large numbers of genes, or one gene weakly co-varies with many transcription factors which may arise when the assayed conditions are inadequately or unevenly sampled. We note that, however, the original CLR is only applied on a symmetric mutual information based matrix while we are dealing with an asymmetric matrix of causality scores. After applying CLR, the network may be still dense and contain spurious edges. Previous studies have shown that the biological networks have some special properties distinct from those of random networks; for example, the network’s out-degree distribution is well approximated by a power law distribution where its in-degree distribution is almost an exponential distribution. Based on those assumptions, we proposed a new regularization method for a directed graph.

The goal of our method is to learn a sparse directed graph from a dense asymmetric causality network (retrieved after applying CLR) satisfying two aforementioned properties. The directed graph’s structure is represented by an indicator matrix denoted by Θ ∈ {0, 1}*^N^*^×^*^N^* where *θ_i,j_* = 1 stands for the existence of edge *i* to *j*, and 0 otherwise. Since the entries are indicators, the in-degree and out-degree of each node in the network can be easily formulated. Specifically, the out-degree of the *i*th node can be represented by 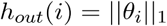 and the in-degree of the *i*th gene is correspondingly represented by 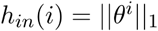, where *θ_i_* and *θ^i^* are the *i*th row and *i*th column of Θ, and *ℓ*_1_ counts the number of nonzero elements since *θ_i,j_* ∈ *j* {0, 1}. Given the asymmetric matrix of causality score *R* with the (*i, j*)th entry as *R_ij_*, the following optimization problem is formulated to learn the structure of the network:

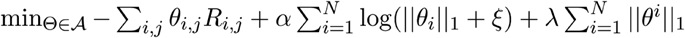

where the feasible set of the network structure is

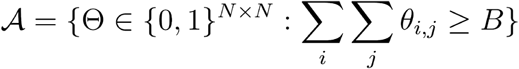

The intuition of the objective function comes directly from the above three assumptions: the first term of the objective is to select the edge with large value of *R_ij_*; the second term is the negative log likelihood of the power law distribution for the out-degree of each gene; the last term is the negative log likelihood of the exponential distribution for the in-degree of each gene. The budget parameter *B* is introduced to prevent trivial solution, and a small positive value *ξ* is used to prevent the numerical issue of log function. The parameter *α* is the exponent of the power law distribution and λ is the parameter of the exponential distribution.

### Benchmarking Scribe with alternative algorithms on inferring causal regulatory network

We follow the same procedure as reported previously (Qiu et al., 2012) to simulate the differentiation of central nervous system (**Eq. 1**), except here we replace the correlated noise in the previous study with independent additive noise for the purpose of simplicity. The data generated through this simulation is regarded as “real-time” dataset.

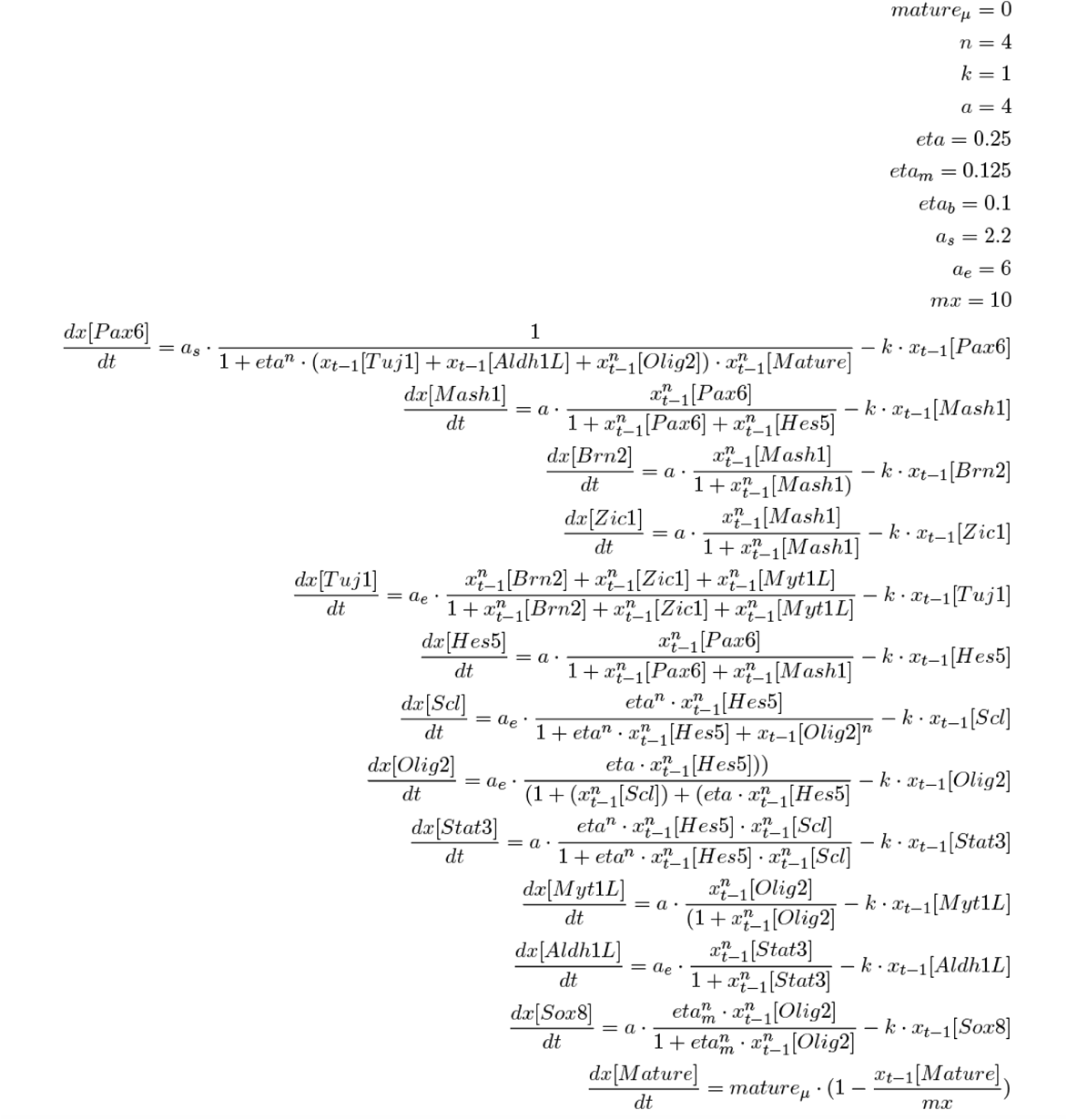

For creating **Fig 1D**, we set the time step as 0.1, samples per simulation as 100, the total number of simulations as 20. We then infer the causal network based on all the 2000 samples using CCM, GC and RDI or uRDI either without conditioning or conditioning on one gene that has the maximal input causality other than the current regulator to the target. Time delay between regulator and target used in all those algorithms is set to be 1. We compare the inferred network with the known network to calculate the AUC (area under curve). The experiment is repeated for 25 times to ensure reliable conclusions. We also increase the standard deviation of the intrinsic noise from 0 to 0.2. ROC (Receiver Operating Characteristic) curve in **Fig 1C** is obtained similarly while setting the simulation based on a linear system where the transition matrix *A* is generated according to the network with non-zero coefficients randomly taken from a uniform distribution *u*(0.75, 1.25). The A matrix is then normalized to *max*(*eigen*(*A*)) * 1.01 to avoid the divergence of the system. The intrinsic noise standard deviation (s.d) is set to be equal to 0.01. All the genes are initialized with a random value *u*(0.5, 2). To infer the causal network, we take 100 samples per simulation and perform the simulation five times, then apply Scribe, CCM and GC on those simulated data points.

To visualize the response, causality and combinatorial regulations as in **Fig 2, Supplementary Figure 2**, a single simulation leading to the neuron fate is used. To create the response and the causality visualization for the two-node motifs (Ma et al., 2009), the network motifs are firstly converted into a set of SDE functions using similar formulations as that used in the above simulation for neuronal differentiation. The expression dynamics is then simulated by setting the initial expression for both genes as 0.01 and followed based on the set of SDE equations **(Fig 3a)**. We used similar procedures to simulate expression of genes under combinatorial regulations with different logic gates and then create the combinatorial regulation visualizations **(Fig 2b)**.

To investigate the importance of temporal coupling and the number of samples on the performance of causal inference, we also simulate three other types of dataset based on the simulated “real time” dataset as following:

1. The RNA-velocity analysis framework estimates both exon and intron expression levels for each cell *i* or *C_i_*. it then calculates the RNA-velocity for each gene (*j*) *V^i^*(*j*) in each cell *i* and predicts the future exon expression of *E^predict^* after Δ*_i_* = 1. Assuming the time delays from all regulators to their putative targets are the same as Δ*t* (or 1), Scribe calculates causality from the potential regulator to the target with the conditional mutual information between the current regulator’s exon expression *x_t_* to the predicted target exon expression *y_t+_*_1_ (or equivalently the estimated RNA velocity value *V_t_*(*y*)) conditioned on the current target exon expression *y_t_* or by the default formula *I*(*x_t_;y_t+_*_1_|*y_t_*) (or alternatively *I*{*x_t_|V_t_*(*y*)*|y_t_*)), Since *x_t_, y_t+_*_1_(*V_t_*(*y*))*, y_t_* are all estimated from the same cell, in theory the gene expression dynamics between *x_t_, y_t+_*_1_*, y_t_* is coupled. To generate RNA-velocity simulation dataset, we randomly select one time point *t* for each cell and collect all genes’ current and the next time point’s expression (*x_i_*(*t*) and *x_i_*(*t* + 1)). RNA velocity for each cell in that time point is then simply calculated as the difference between next time point and current time point’s gene expression (*V_t_*(*y*) *= X_i_*(*t +* 1) *− x_i_*(*t*)).
2. To generate live-imaging simulation dataset, we first randomly select 13 cells where for each cell, a different gene is chosen and is followed over the entire developmental process.
3. To generate pseudotime dataset, similar to RNA-velocity, we randomly select one time point *t* for each cell and collect all genes’ expression at that time point. Then all data points from each cell at different time point is pooled and used as input to Monocle 2 for trajectory inference, we then set the beginning of the simulation as root state for the trajectory and order cells based on the inferred pseudotime to form a pseudotime series.

To create **Fig 6B** or **Supplementary Figure 6A**, five replicates each with 2000 data points are used for each algorithm. For **Fig 6C** and **Supplementary Figure 6B**, the same analysis is performed but with data downsampled to 200, 400, 600, 800, 1000, 1200, 1400, 1600, 1800 or 2000 data points.

### Details on analyzing datasets used in this study

**Inferring causal network with pseudotime ordered scRNA-seq datasets**. Lung data is processed as described previously (Qiu et al., 2017a). Categorization of pneumocyte specification markers into either early and late groups used for benchmarking is based on references(Qiu et al., 2017a; Treutlein et al., 2014).

The LPS data was pre-processed as described previously (Qiu et al., 2017b) while the trajectory is reconstructed with the reversed graph embedding (Qiu et al., 2017a) on the same set of ordering genes used in this study. Only the path with wild-type cells is used for causal network inference. Regulators and targets, and the regulatory network used for benchmarking are collected from references (Amit et al., 2009) and reference (Garber et al., 2012), respectively.

Olsson data is processed as described previously. The master regulators, transcription factors and downstream targets, and the regulatory network used for benchmarking are collected from reference (Qiu et al., 2017a) and references (Su et al., 2017), respectively.

Paul data is processed as described previously. Only the path leading to the erythrocytic fate is used for reconstructing the causal regulatory network. The regulatory network responsible for the differentiation of erythrocyte cells used for benchmarking is collected from (Swiers et al., 2006).

**Infer causal network with RNA-velocity**. The data of the chromaffin cell “RNA-velocity” analysis is retrieved from (http://pklab.med.harvard.edu/velocyto/notebooks/R/chromaffin.nb.htmn. We use the estimated exon expression to reconstruct the trajectory for the chromaffin cell commitment. Only cells on the path from the Schwann cell progenitors to mature chromaffin cells are used to infer the casual network. Two different formulations, *^I^*(*^x^f, yt+i\yt*) (or *I*(*^x^t',Vt*(*y*)*\yt*))*_t_* can be used to infer causal networks with data from RNA-velocity. In this study, we apply the first formulation.

**Inferring causal network with live-image data**. Lineage-resolved live-imaging data for *C. elegans* early embryogenesis is obtained from Waterston lab. Raw fluorescence intensity signal is directly used for causal network inference. We note two caveats in analyzing the reporter data with Scribe. First, although the promoter-fusion data sheds light on the induction kinetics of the TF of interest, once the fluorescent reporter is expressed it follows the trafficking and degradation kinetics of the histone protein, and not the TF. Second, the time series for each TF was captured in a different embryo, so this may introduce noise that obscures the regulator/target relationships between the TFs although the *C. elegans* development process is highly robust. Nevertheless, this data set represents an unprecedented view of TF activity at high spatiotemporal resolution during the early development of a complex organism.

